# Structural Analysis of Receptor Binding Domain Mutations in SARS-CoV-2 Variants of Concern that Modulate ACE2 and Antibody Binding

**DOI:** 10.1101/2021.08.25.457711

**Authors:** Dhiraj Mannar, James W Saville, Xing Zhu, Shanti S. Srivastava, Alison M. Berezuk, Steven Zhou, Katharine S. Tuttle, Andrew Kim, Wei Li, Dimiter S. Dimitrov, Sriram Subramaniam

**Author notes:** These authors contributed equally (Dhiraj Mannar, James W Saville).

## Abstract

The recently emerged SARS-CoV-2 South African (B. 1.351) and Brazil/Japan (P.1) variants of concern (VoCs) include a key mutation (N501Y) found in the UK variant that enhances affinity of the spike protein for its receptor, ACE2. Additional mutations are found in these variants at residues 417 and 484 that appear to promote antibody evasion. In contrast, the Californian VoCs (B.1.427/429) lack the N501Y mutation, yet exhibit antibody evasion. We engineered spike proteins to express these RBD VoC mutations either in isolation, or in different combinations, and analyzed the effects using biochemical assays and cryo-EM structural analyses. Overall, our findings suggest that the emergence of new SARS-CoV-2 variant spikes can be rationalized as the result of mutations that confer either increased ACE2 affinity, increased antibody evasion, or both, providing a framework to dissect the molecular factors that drive VoC evolution.

## Introduction

Recent genomic surveillance efforts tracking the global spread of SARS-CoV-2 have identified the emergence and rapid spread of several variants. Variants B. 1.1.7 & VOC 202102/02 (“UK”), B.1.351 (“South Africa”), P.1 (“Japan/Brazil”) and B.1.427 & B.1.429 (“California”) have all been identified by the American Centers for Disease Control and Prevention (CDC) as variants of concern (VoCs)(CDC, 2021a). VoCs are designated as having evidence demonstrating increased transmissibility, increased disease severity and/or a significant impact on diagnostics, treatments and vaccines (CDC, 2021a, 2021b; FDA, 2021; Moore and Offit, 2021; Wang et al., 2021d). Common amongst these six VoCs are mutations within the receptor-binding domain (RBD) in the spike glycoprotein (S protein). The S protein protrudes from the surface of the virus and facilitates viral attachment, fusion and entry into cells via its binding partner human angiotensin-converting enzyme 2 (ACE2) (Shang et al., 2020). Additionally, the S protein is the major target of the humoral immune response, with the majority of currently developed vaccines using this protein as their major antigenic component (Krammer, 2020). The RBD within the S protein constitutes the region against which the majority of neutralizing antibodies are directed, highlighting the importance of this region for viral infection and antibody neutralization (Barnes et al., 2020a; McCallum et al., 2021; Piccoli et al., 2020).

Mutations within the RBD of the SARS-CoV-2 S protein may confer enhanced viral fitness by increasing the affinity of the S protein for ACE2 and/or decreasing the neutralization activity of antibodies produced by the humoral response. Additionally, RBD mutations may allow for increased expression or presentation of the S protein on the viral surface. Figure 1B and C show the S protein mutations present in each of the six VoCs, with the majority of common mutations found within the RBD. Additionally, three out of the four VoC RBD mutations (L452R, E484K, N501Y) are located within the receptor-binding motif (RBM) which comprises the interaction interface between the S protein and ACE2. The one RBD mutation occurring outside of the RBM - K417N/T - additionally exhibits ambiguity in mutation; with the P.1 strain mutated to threonine (K417T) and the B. 1.351 strain mutated to asparagine (K417N) (Figure 1B). In the present study, we aim to understand the individual and combinatorial contributions that each of these common VoC RBD mutations has on enhancing aspects of viral fitness.

**Figure 1.**
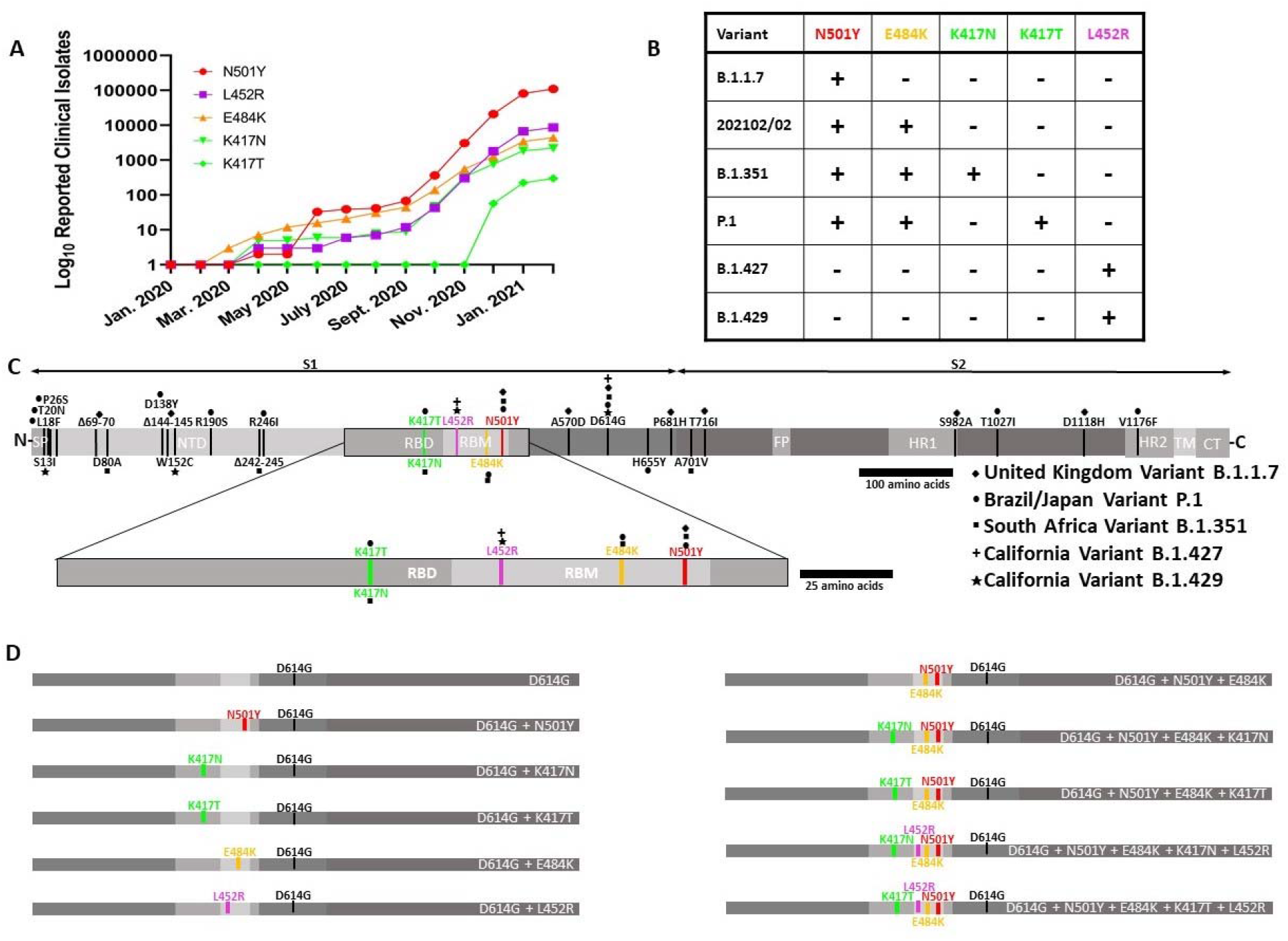
The global prevalence of SARS-CoV-2 VoC RBD mutations and their locations within the S protein. **(A)** Global occurrences of each VoC RBD mutation over time, computed using the sum of clinical isolate entries each month deposited into the GISAID database (https://www.gisaid.org/). **(B)** Summary of the RBD mutations present in each VoC. **(C)** SARS-CoV-2 spike glycoprotein amino acid open reading frame (ORF) with VoC mutations indicated. An expanded portion of the RBD is provided to highlight the common RBD mutations between the VoCs. Relevant features are indicated: SP (Signal Peptide); NTD (N-terminal Domain); RBD (Receptor Binding Domain); RBM (Receptor Binding Motif); FP (Fusion Peptide); HR1 (Heptad Repeat 1); HR2 (Heptad Repeat 2); TM (Transmembrane Domain); CT (Cytoplasm Domain). **(D)** Summary of the SARS-CoV-2 spike glycoprotein constructs employed in this study. VoC RBD mutations were expressed in isolation, in naturally occurring combinations, and in novel combinations to assess the relative individual and combinatorial effects of these mutations. All constructs contain the D614G mutation as background and this was defined as the wild-type construct throughout the study.

Using eleven S proteins with different complements of mutations, we systematically dissect the contributions of the VoC RBD mutations towards increasing ACE2 affinity and evading neutralizing antibodies (Figure 1D) using cryogenic electron microscopy (cryo-EM) structural analyses and assays that measure ACE2 and antibody binding. We also constructed novel, and as yet unreported combinations of RBD mutations to explore the properties of variants that may emerge in the future as the SARS-CoV-2 strains evolve. Our analyses make use of trimeric HexaPro stabilized S protein ectodomain constructs which differ from native S protein trimers by the addition of six stabilizing proline mutations (F817P, A892P, A899P, A942P, K968P, V969P) and the transmembrane domain replaced with a trimerization motif (Hsieh et al., 2020). The HexaPro construct produces substantially higher yields in mammalian cell culture and has increased thermostability which facilitates the structural and biochemical experiments presented here.

## Results

### The N501Y, E484K and L452R mutations drive increased S protein - ACE2 binding affinity

To investigate the effects of VoC RBD mutations on ACE2 binding, we expressed and purified recombinant spike ectodomain proteins bearing VoC RBD mutations in isolation and combination (Figure S1), which we utilized in biolayer interferometry (BLI) experiments (Figures 2A-B, S2). When compared to wild-type (D614G), spikes harbouring combinations of RBD mutations found in circulating VoCs exhibited increased ACE2 binding affinities. The individual addition of N501Y, E484K or L452R mutations increased ACE2 binding affinity, and the increased affinity conferred by the N501Y and E484K mutations in isolation were preserved in combination in the D614G + N501Y + E484K construct, yielding the highest affinity ACE2 binder. Mutations at the 417 position (K417N/T) decreased the affinity for ACE2 both in isolation (D614G + K417N/T) and when introduced into the D614G + N501Y + E484K construct. Interestingly, the K417N mutation reduced ACE2 affinity to a higher extent than the K417T mutation (both in isolation and when combined with D614G + N510Y + E484K). Taken together, these results demonstrate that the amalgamation of spike RBD mutations present in circulating VoCs enables increased ACE2 affinity, which is mainly driven by N501Y (B.1.1.7), L452R (B.1.427/B. 1.429), and the combinatorial effect of both N501Y, and E484K (P.1, B.1.351, VOC 202102/02).

**Figure 2.**
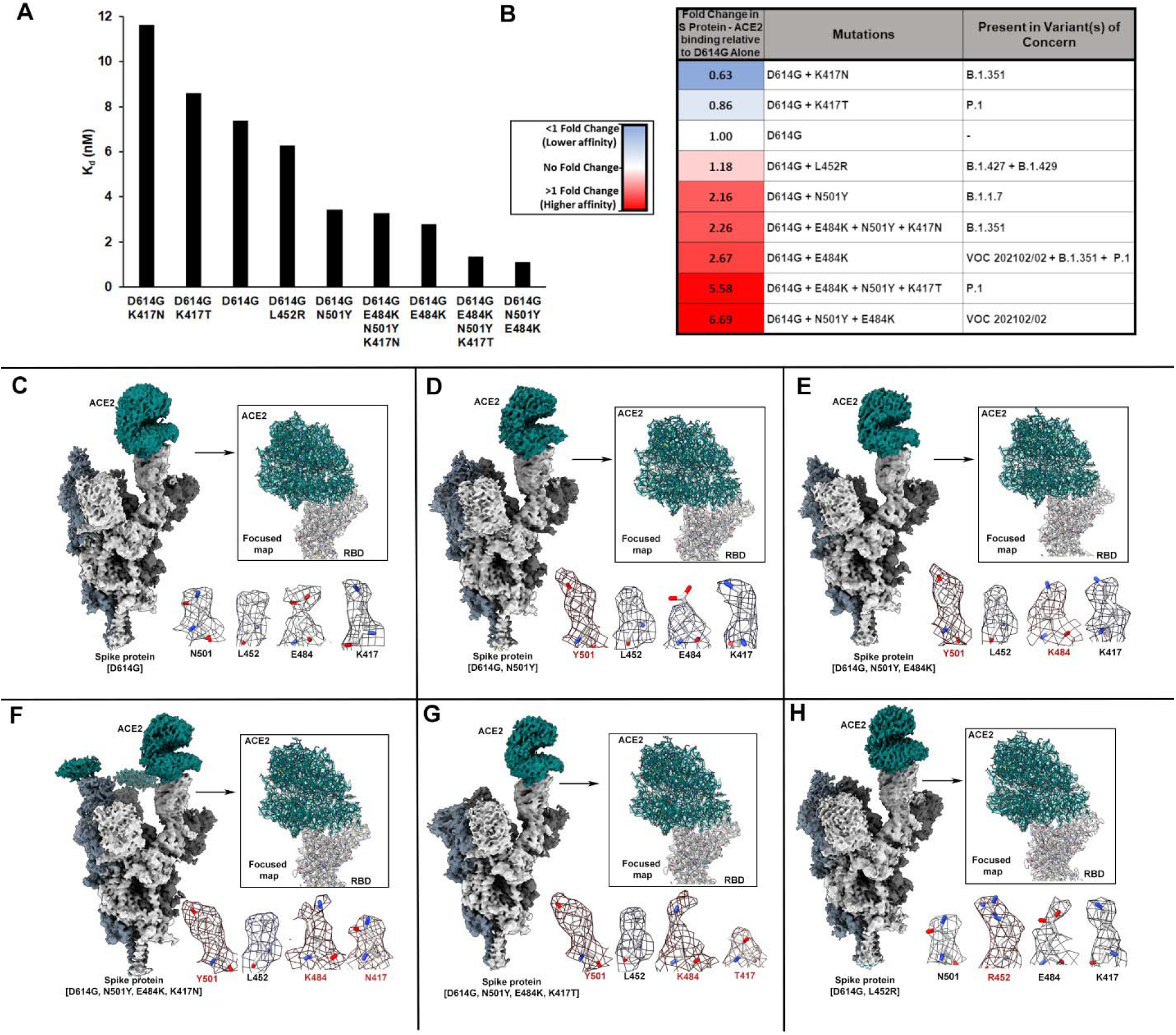
Complete sets of VoC RBD mutations increase S protein trimer - ACE2 binding affinity. **(A)** Affinity (K_d_) measurements for VoC RBD mutant S protein - ACE2 binding as measured by biolayer interferometry (BLI). **(B)** Relative fold change differences in S protein - ACE2 affinity (K_d_) relative to D614G alone. **(C-H)** Structures of VoC spike-ACE2 complexes characterized in this study. Shown for each complex studied are density maps for the overall complex at the end of global structure refinements, improved focused density maps at the ACE2-RBD contact zones, and visualization of densities at mutational positions within each variant spike. Densities at sites harbouring mutations are highlighted with red text.

### Mutational effects on ACE2 binding are mediated by subtle side-chain rearrangements at the S protein – ACE2 interface

To understand the structural effects of the various VoC RBD mutations on ACE2 binding we conducted cryo-EM studies on unbound spike trimers and ACE2–spike complexes (Figure 2C-H, S3-S14, Table S1). Resulting structures were obtained at average resolutions of ~2.3-3 Å (Figures S3-S14, Table S1). The *apo*- and ACE2-complexed S protein structures show no significant global changes in secondary or quaternary structure as a result of the various mutations when compared to D614G (Figure S15). However, focused refinement of the S protein – ACE2 interface revealed side chain rearrangements that may account for the observed differences in binding affinity as outlined below:

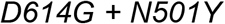

The cryo-EM structure of ACE2 bound to the D614G + N501Y mutant spike (Figures 3B, S16B) shows the same features at the RBD-ACE2 interface as in our previously reported structure of the N501Y-ACE2 complex in the absence of the D614G mutation (Figure 3A, S16A). Y501 in the spike protein and Y41 in the ACE2 receptor engage in a perpendicular y-shaped π–π stacking interaction (Zhu et al., 2021). Additionally, superposition of the RBD in all RBD-ACE2 structures reveals a ~2.4 Å displacement of an ACE2 helix distal to the RBD binding helix when compared to complexes with N and Y at residue 501 reflecting the impact of this change (Figure S16C).

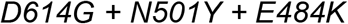

**Figure 3.**
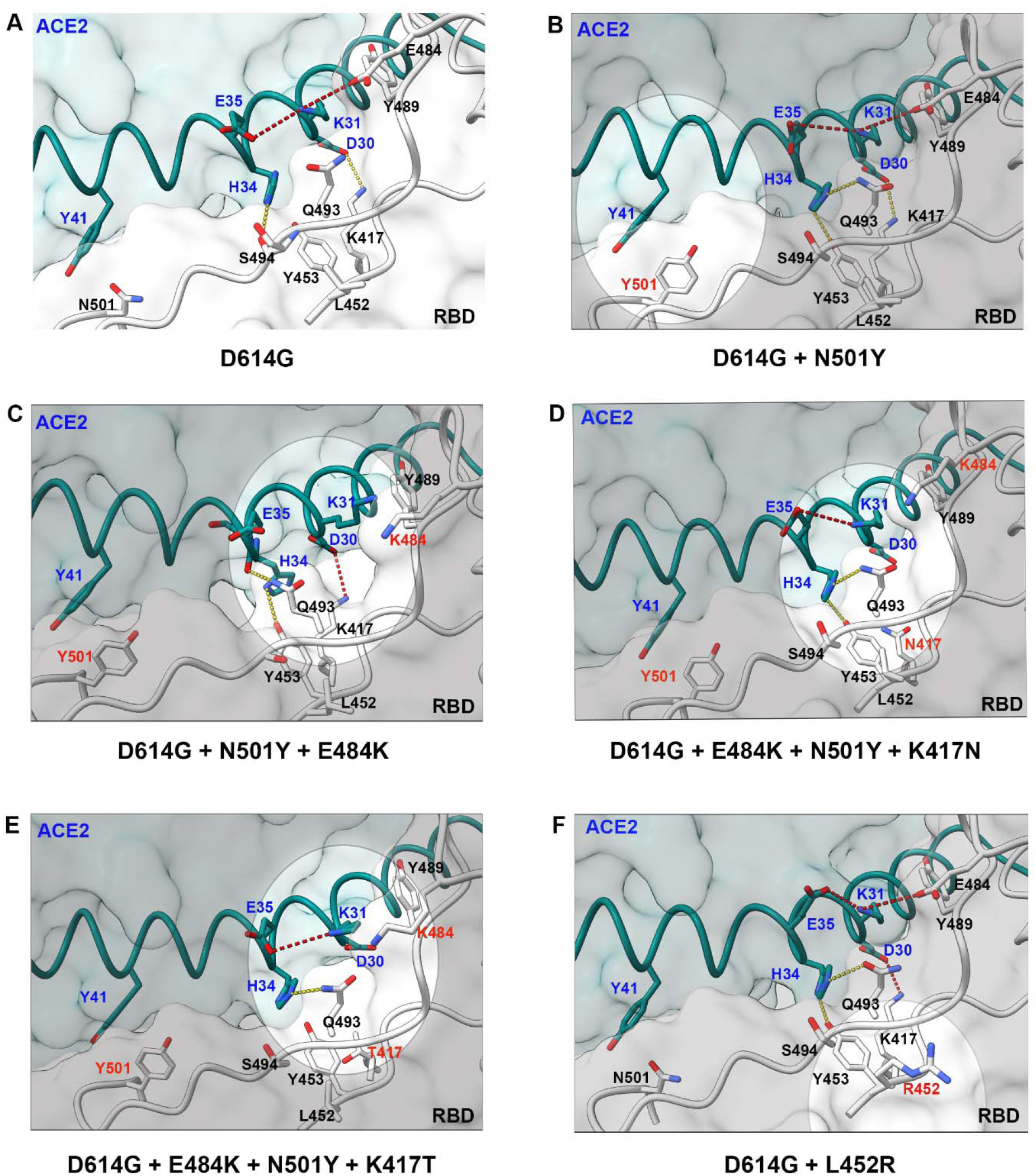
CryoEM structures of wild-type and VoC RBD-ACE2 interfaces. **(A-F)** Zoomed in views of the RBD-ACE2 binding interfaces for the six S protein – ACE2 structures. Focused refinement of the RBD-ACE2 interface reveals distinct S protein and ACE2 side-chain rotamer arrangements for the various VoCs. Mutated residues are labeled in red and adjacent residues of interest are highlighted within ovals. ACE2 residues are labeled in blue while RBD residues are labeled in black. Hydrogen bonds and electrostatic interactions are shown as yellow and red dotted lines, respectively. Oxygen and nitrogen heteroatoms are colourized in red and blue respectively.

Analysis of the D614G + N501Y + E484K mutant spike in complex with ACE2 reveals local rearrangements resulting in unambiguous rotamer placement of both H34 within ACE2 and Q493 within the spike RBD (Figures 3C, S17). The resulting H34 rotamer yields space which accommodates an alternate Q493 rotamer closer to ACE2 relative to the D614G spike, allowing it to be positioned within hydrogen-bonding distance of the main chain carbonyl of H34. Additionally, the positioning of K31 within ACE2 is shifted relative to the D614G spike, adopting a position within pi-cation bonding distance to Y489 within the RBD (Figure S17C). K484 extends parallel to the RBD-ACE2 plane of interaction, likely due to electronic repulsion from K31, and adopts a position ~7.5 Å from E35. These subtle changes in intermolecular interactions enabled upon H34 repositioning suggest a basis for the enhanced ACE2 affinity observed for the D614G + N501Y + E484K mutant spike relative to D614G + N501Y.

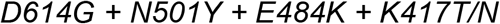

The mutation of K417 to T or N resulted in loss of the K417 - D30 salt bridge within the ACE2-spike complex, providing a basis for the decreased ACE2 binding affinities conferred by these two mutations (Figures 3D-E, S17D-E). In contrast to the D614G + N501Y + E484K - ACE2 complex, H34 rotamer placement is ambiguous within these complexes, with the predominant densities corresponding to H34 facing toward the K484 interface. Additionally, Q493 adopts a rotamer which faces away from ACE2, and K31 is positioned to face both H34 and Q493.

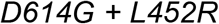

Structural comparison of D614G-ACE2 and D614G+L452R-ACE2 complexes reveals no significant changes at the RBD-ACE2 interface (Figure 3F, S17F), indicating the enhanced ACE2 affinity afforded by L452R is not due to modulation of direct ACE2 contacts. In contrast to L452, the side chain of R452 extends farther away from the RBD core (Figure S18A) and is better exposed to solvent, suggesting that R452 may enhance the solvation of the RBD in the up position. In addition to solvation effects, the L452R substitution introduces a positive charge at position 452 which may increase the electrostatic complementarity between the RBD and ACE2. Figure S18B shows the increase in electropositivity at position 452 upon L452R substitution, with position 452 approximately 13 Å away from the highly electronegative site on ACE2 centered at E35. Thus, in contrast to the local rearrangements observed at the RBD-ACE2 interface for the N501Y, E484K and K417N/T mutations, the binding effect of the L452R mutation is likely mediated by solvation and/or electrostatic complementarity effects.

### Mutations E484K, L452R and K417N/T facilitate decreased antibody binding

We next sought to evaluate the effect of VoC RBD mutations on antibody binding. We selected a panel of previously reported antibodies which cover the four distinct anti-RBD antibody classes (Barnes et al., 2020b, Table S3) and an ultrapotent antibody, S2M11, that uniquely binds two neighbouring RBDs (Tortorici et al., 2020). Antibody binding was quantified via enzyme-linked immunosorbent assay (ELISA) (Figures 4, S19). As expected, class 3 (S309) and class 4 (CR3022) antibodies, whose footprints did not span VoC mutations, exhibited relatively unchanged binding across all variant spikes (Figure 4B). Mutations at position 417 of the S protein to either N or T abolished or significantly reduced ab1 (Li et al., 2020a) (class 1 like) binding respectively, demonstrating the importance of K417 within the molecular epitope of ab1. Similarly, the E484K mutation resulted in loss of binding to ab8 (Li et al., 2020b) (class 2) and S2M11, highlighting the critical nature of E484 within the epitopes of these antibodies. L452 sits peripherally within the footprint of S2M11 and mutation of this residue to R452 reduced but did not abolish its binding, possibly via steric or charge mediated effects, or by allosteric modulation of direct contacts. Taken together, these results suggest the escape of antibody binding from the four major anti-RBD classes is primarily mediated by modulation of direct contacts at mutational sites.

**Figure 4.**
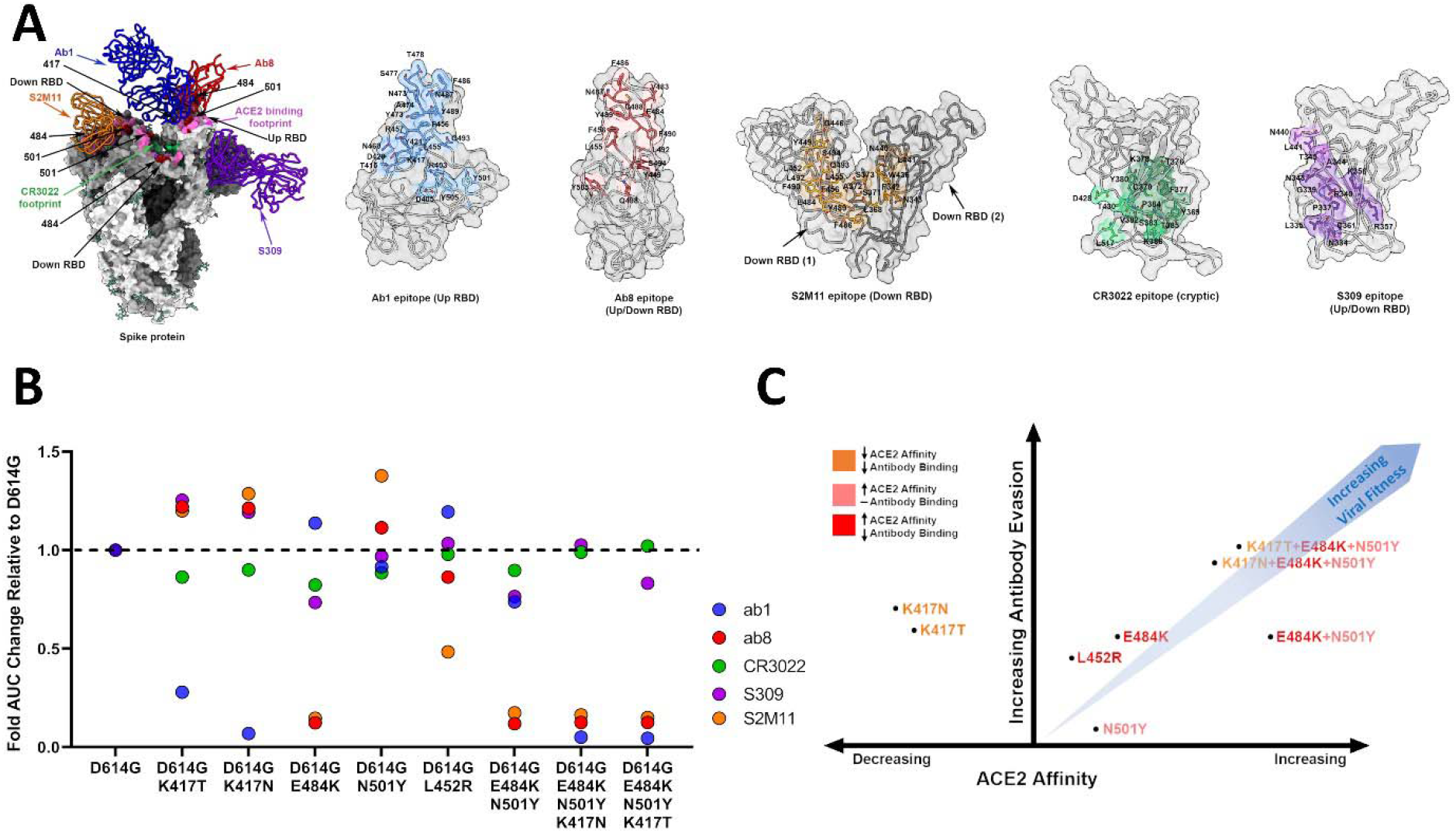
Monoclonal antibody binding against SARS-CoV-2 S proteins containing VoC RBD mutations. **(A)** Mapping of Ab1, Ab8, CR3022, S309 and S2M11 antibody footprints onto SARS-CoV-2 trimers and RBDs. Direct amino acid contacts for each individual antibody footprint are highlighted separately. **(B)** Area under the curve (AUC) fold changes in ELISA binding assays relative to D614G alone for Ab1, Ab8, CR3022, S309, and S2M11. **(C)** Qualitative 2D plot describing VoC RBD mutational effects on ACE2 and antibody binding. The mutations are grouped into three colour categories: Orange, mutations that decrease ACE2 affinity and antibody binding; Pink, mutations that increase ACE2 affinity and do not significantly impact antibody binding; and Red, mutations that increase ACE2 affinity and decrease antibody binding. The blue arrow shows the theoretical trend towards increasing viral fitness, however its slope should not be interpreted as an exact, quantitatively determined measure.

### Novel RBD mutant combinations preserve but do not enhance effects on ACE2 affinities and antibody binding

Having determined that all full complements of VoC RBD mutations results in increased ACE2 binding and various extents of antibody evasion, we aimed to assess the functional effects of novel VoC RBD mutational combinations that have not yet been reported, but represent combinations of mutations already observed. VoCs harbouring N501Y exhibit a spectrum of additional RBD mutations (B.1.1.7: N501Y, VOC 202102/02: E484K, N501Y, B.1.351: E484K, N501Y, K417N and P.1: E484K, N501Y, K417T), while VoCs containing L452R (B.1.427/B.1.429) seemingly exclude N501Y, K417N/T, and E484K mutations, though there has been a recent report of E484Q co-mutation with L452R in India (B.1.617.1). In order to assess if such patterns of evolution are due to incompatibility of these mutations, we constructed and expressed recombinant spike ectodomains combining L452R with the full complement of either B.1.351 and P.1 RBD mutations, and evaluated ACE2 and antibody binding of these mutants (Figure S20 A,B). Neither of these novel combinations conferred enhanced ACE2 affinities when compared to B.1.351 and P.1 RBD mutant spikes. Notably, both novel combinatorial mutants still exhibited enhanced ACE2 binding when compared to wild type (Figure S20B). The addition of L452R to both constructs preserved the antibody evasive properties for K417N/T against ab1 and E484K against both ab8 and S2M11 (Figure S20C). These results indicate that while the L452R mutation is not mutually exclusive with the complement of RBD mutations in B.1.351 and P.1 variants with regards to reduction of neutralizing antibody binding, the increase in ACE2 binding affinity conferred by the L452R mutation in isolation (Figure 2A-B), is absent when combined with B.1.351 and P.1 RBD mutations.

## Discussion

We have dissected the relative roles of circulating VoC RBD mutations with regards to both ACE2 affinity and antibody binding (Figure 4C). Our results demonstrate that individual mutations may be classified as resulting in either (a) increased RBD-ACE2 affinity alone (N501Y), (b) reduced ACE2 affinity and reduced antibody binding (K417N/T), or (c) a simultaneous increase in ACE2 affinity and reduced antibody binding (E484K, L452R). These individual effects are preserved when mutations are combined to reflect full complements of VoC RBD mutations, demonstrating their modular nature. Furthermore, these results suggest that RBD evolution follows a trajectory aimed towards simultaneous enhancement of receptor affinity and reduction of neutralizing antibody binding. It is noteworthy that all VoCs containing K417N/T mutations also contain the N501Y and E484K mutations. Given that K417N/T mutations serve to diminish antibody binding at a cost to ACE2 affinity, the conditional presence of ACE2 affinity enhancing mutations may represent a compensatory mutational mechanism. Consistent with this hypothesis, analysis of deposited spike sequences in the GISAID database reveals that K417N/T mutations do not occur independently of N501Y and E484K mutations (Figure S21). In contrast, K417N/T mutations are not a prerequisite for the occurrence of mutations which increase ACE2 affinity (N501Y) or simultaneously increase ACE2 affinity and decrease antibody binding (E484K, L452R) (Figure S21).

Our described effects on ACE2 binding and antibody evasion imparted by VoC RBD mutations are in agreement with recent reports (Chen et al., 2021; Collier et al., 2021; Dejnirattisai et al., 2021; Laffeber et al., 2021; Liu et al., 2021; Wang et al., 2021a, 2021b; Wibmer et al., 2021; Yuan et al., 2021), with the exception of the enhanced ACE2 affinity conferred by E484K. Several studies have reported conflicting data surrounding the effect of E484K on ACE2 binding where both decreased (Upadhyay et al., 2021; Wang et al., 2021c; Yuan et al., 2021) and increased (Laffeber et al., 2021; Tanaka et al., 2021) affinities are observed. A variety of biophysical techniques, spike protein domains, and ACE2 constructs were employed across these studies which could account for the contrasting results. Most importantly, the E484K mutation was amongst several mutations selected via *in vitro* evolution to affinity maturate the RBD for enhanced ACE2 binding (Zahradník et al., 2021), demonstrating a clear role for increasing ACE2 binding affinity.

The present study highlights the potential for antibody evasion by VoC RBD mutations via antibody binding assays using a panel of monoclonal antibodies. To estimate the potential effect of VoC RBD mutations on RBD binding by naturally acquired antibodies during SARS-CoV-2 infection, we selected PDB entries of SARS-CoV-2 spike or RBD complexes with antibody fragments isolated from convalescent patients (Table S2). Using this structural data, we evaluated the frequency of positions corresponding to RBD mutations in VoCs within the footprint of 27 selected antibodies. The majority of deposited human-derived neutralizing antibodies bound the RBD with footprints spanning at least one of the positions corresponding to RBD mutation in VoCs (Figure S22B). Of these antibodies, the majority interacted with more than one position corresponding to RBD mutations in VoCs (Figure S22C). Of the VoCs, B.1.351, P.1 and VOC 202102/02 possess mutations that are collectively recognized by the majority of the antibodies selected, suggesting these VoCs may exhibit the greatest RBD-directed antibody escape during human infection (Figure S22D).

We additionally generated novel combinations of RBD mutations by introducing L452R into B.1.351 and P.1 constructs. While these mutational combinations enable enhanced ACE2 binding compared to wild-type spikes, the increase in ACE2 binding affinity conferred by the L452R mutation in isolation was not preserved. We demonstrate that these novel constructs retain antibody evasive properties when tested for antibody binding using a panel of monoclonal antibodies. Although there are many factors governing viral evolution, these results suggest that the independent evolution of L452R bearing spikes and N501Y, K417N/T, and E484K bearing spikes may be explained due to a lack of synergistic increase in ACE2 binding upon combination of these mutations. Such combinations may however still evolve in the future as a result of increased antibody escape.

The cryo-EM structures of all 5 VoC RBD-mutated spike trimers, both in isolation and in complex with ACE2, provide insights regarding the molecular basis for observed changes in ACE2 affinities. The combination of enhanced intermolecular interactions due to the concomitant repositioning of H34 and Q493 in the D614G + N501Y + E484K - ACE2 complex provides structural rationale for the increased ACE2 binding affinity relative to the D614G + N501Y spike. Although hydrogen bonding with Y453 is possible in both H34 rotamers (Figure S17), the dominant rotamer positioning of H34 in the D614G + N501Y + E484K - ACE2 complex enables it to participate in additional favourable intermolecular interactions with Y453 (hydrogen bond +OH/ π) and L455 (CH/π), yielding estimated interaction energies of −10.29 and −2.75 kcal/mol, respectively (Watanabe et al., 2021). The mechanism of H34 rotamer stabilization in response to the E484K mutation remains unclear at present, although the repositioning of Q493 in this structure permits the formation of an intermolecular hydrogen bond with the main chain carbonyl of H34. This is in contrast to all other structures of spike protein-ACE2 complexes in which Q493 is positioned in close proximity to the main chain RBD carbonyls of F490 and L492, and is well poised to participate in intramolecular hydrogen bonds (Figure S17). Finally, the positioning of K31 within pi-cation bonding distance to Y489 in this structure suggests an additional intermolecular interaction that may enhance ACE2 affinity (Figure S17C). It should be noted that although the intermolecular K31-E484 salt bridge is lost upon inclusion of the E484K mutation, this electrostatic interaction is likely intramolecularly distributed between ACE2 residues E35 and K31, thus limiting the contribution of the K31-E484 interaction with regards to ACE2-RBD binding. The positioning of H34 away from residue 484 in all RBD-ACE2 crystal structures reported (PDB: 6M0J, 6VW1, 7NXC) agrees with our assessment that this represents the more energetically favourable rotamer with regards to the stability of the RBD-ACE2 complex. This structural basis is consistent with previous reports implicating H34 as a major contributor to the SARS-CoV-2 RBD-ACE2 interaction (Glasgow et al., 2020) and reports demonstrating the H34A mutation in ACE2 enhances SARS-CoV-2 spike binding (Chan et al., 2020; Tanaka et al., 2021). This is likely due to closer positioning and flexibility of RBD resides such as Q493. Recent reports have suggested that the E484K mutation may enhance ACE2 binding via increasing electrostatic complementarity between ACE2 and the RBD (Dejnirattisai et al., 2021; Fantini et al., 2021), and the structures reported here are consistent with that hypothesis. The structural basis for the observed discrepancies in ACE2 binding between the K417T and K417N mutations remains unclear.

Since L452 is distal to the ACE2-RBD interface, it has been previously suggested that the L452R mutation may increase ACE2 affinity via allosteric modulation of the residues promoting the RBD-ACE2 interaction (Deng et al., 2021), or via electrostatic effects (Motozono et al., 2021). We did not observe any allosteric changes in our structures, rather we highlight the enhanced RBD-ACE2 electrostatic complementarity and potentially increased RBD solvation as explanations for the increased ACE2 affinity conferred by R452. Protein-protein interaction studies have predicted that long-range electrostatic complementarity plays a role in determining complex association rates (Schreiber et al., 2009). Therefore, the increased electrostatic complementarity between ACE2 and the RBD due to R452 may enhance ACE2 affinity via increasing the probability of forming favourable RBD-ACE2 binding orientations. The increased solvation and electrostatic complementarity explanations are not mutually exclusive and may contribute to increased ACE2 affinity in combination.

Although we have focused the present study on RBD mutations present within VoCs, it is possible that mutations elsewhere in the S protein (particularly in the N-terminal domain) also play a significant role in antibody evasion and may affect ACE2 binding (McCallum et al., 2021). Mutations outside of the S protein open reading frame may additionally contribute towards increased viral fitness. Future studies will likely provide further insights into these aspects of SARS-CoV-2 infection and global spread.

## Acknowledgments and Funding

This work was supported by awards to S.S. from a Canada Excellence Research Chair Award, the VGH Foundation, and Genome BC, Canada and a generous donation from the Tai Hung Fai Charitable Foundation. W.L. and D.S.D. were supported by the University of Pittsburgh Medical Center (UPMC). D.M. is supported by a CIHR Frederick Banting and Charles Best Canada Graduate Scholarship Master’s Award (CGS-M). J.W.S is supported by a CIHR Frederick Banting and Charles Best Canada Graduate Scholarships Doctoral Award (CGS-D) and a UBC President’s Academic Excellence Initiative PhD Award. We thank Sagar Chittori for assistance with microscopy screening and collection.

## Author contributions

J.W.S and D.M. performed the molecular cloning. J.W.S., D.M., S.Z., S.S.S., carried out expression and purification of the spike proteins and antibody fragments. A.K. performed the BLI binding assays under the supervision of W.L. and D.S.D.. D.M. and J.W.S. performed the antibody binding experiments. A.M.B., and K.S.T. collectively carried out the experimental components of cryo-EM and electron microscopy including specimen preparation and data collection. X.Z. carried out all computational aspects of image processing and structure determination. D.M., X.Z., S.S.S. and S.S. interpreted and analyzed the cryo-EM structures. A.K., W.L. and D.S.D. provided the plasmids for VH-ab8 and IgG ab1 as part of a collaboration between the Subramaniam and Dimitrov laboratories on SARS-CoV-2. D.M., J.W.S., S.S.S., and S.S. drafted the initial manuscript with input from all authors.

## Data and Materials Availability

The density maps and atomic coordinates for the structures reported will be made publicly available in the Electron Microscopy Data Bank.

## Materials and Methods

### Key Resources Table

#### Kits and reagents

|  |  |  |
| --- | --- | --- |
| Pierce 1-Step Ultra Substrate Solution | ThermoFisher | Cat. # 34028 |
| Q5 Site-Directed mutagenesis kit | NEB | Cat. # E0554S |

#### Antibodies

|  |  |  |
| --- | --- | --- |
| VH ab8 | (Li et al. 2020) | N/A |
| IgG1 ab1 | (Li et al. 2020) | N/A |
| IgG1 CR3022 | (Yuan et al. 2020) | N/A |
| Fab S309 | (Pinto et al. 2020) | N/A |
| Fab S2M11 | (Tortorici et al. 2020) | N/A |
| goat anti-human IgG | Jackson ImmunoResearch | Cat. # 109-035-088 |

#### Recombinant proteins

|  |  |  |
| --- | --- | --- |
| Ace2 (18-615) | New England Biolabs | Cat. # 73775S |

#### Cell lines

|  |  |  |
| --- | --- | --- |
| Expi293F | ThermoFisher | Cat# A14527 |

#### Recombinant DNA and Oligonucleotides

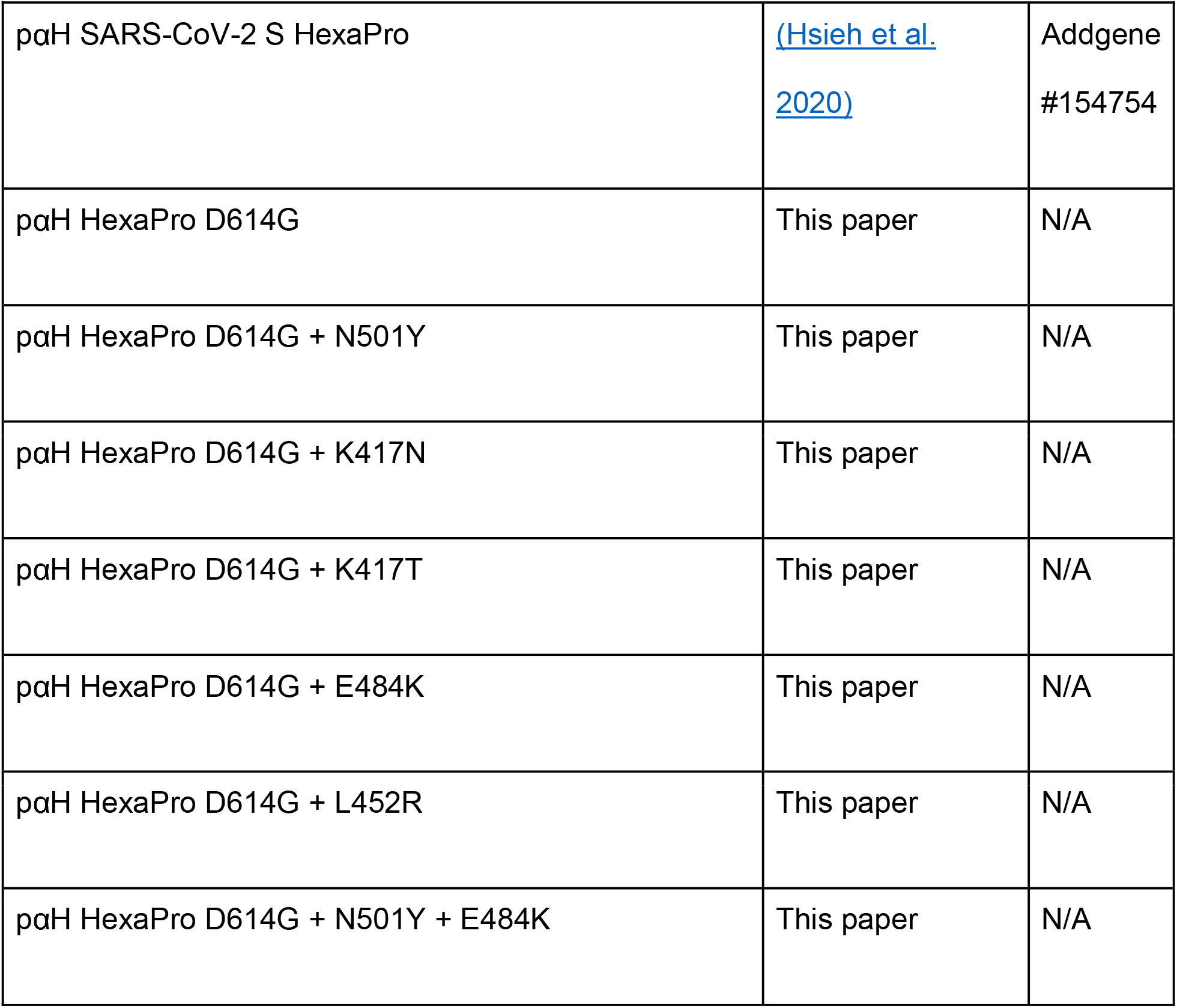
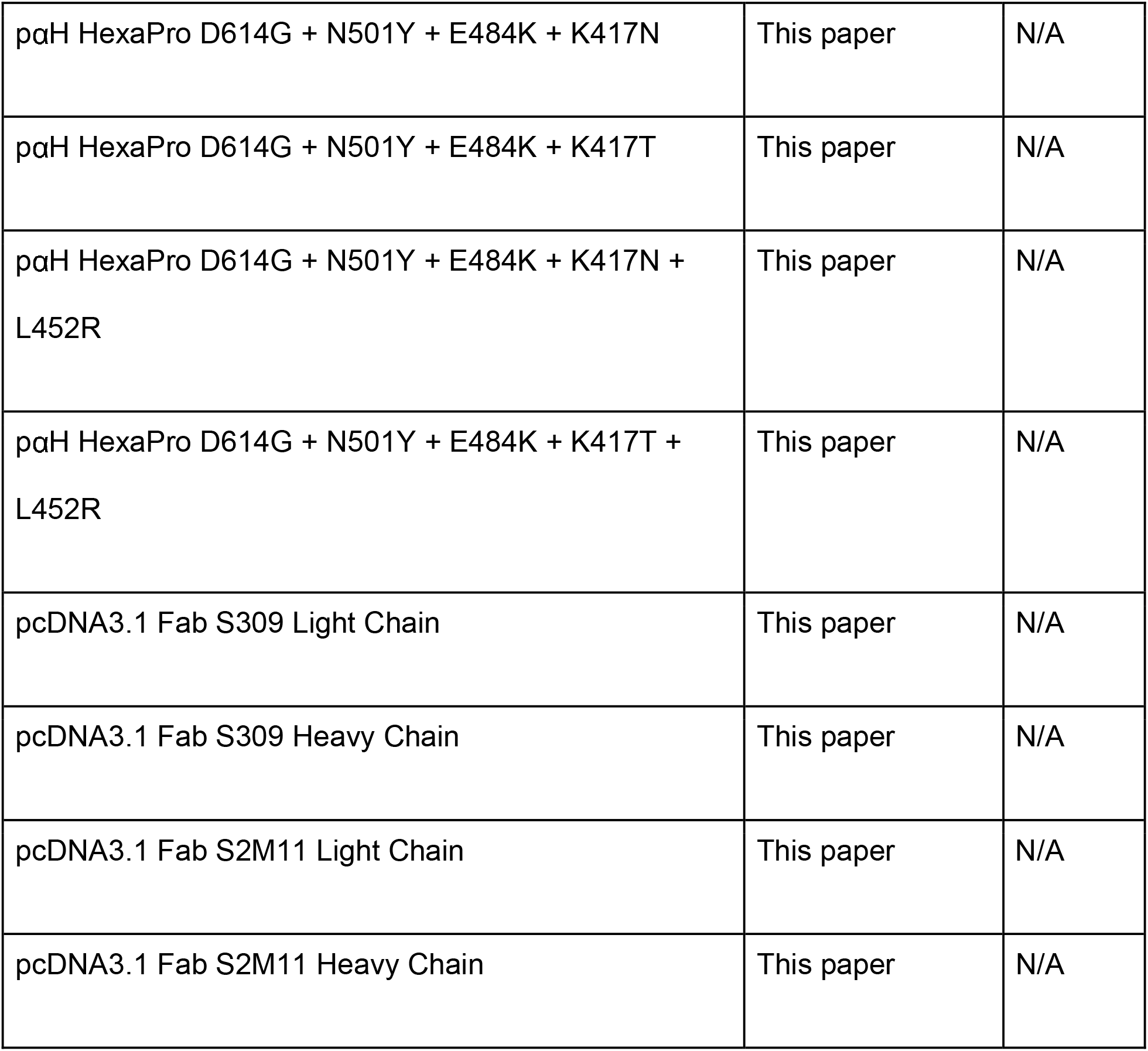

#### Software and Algorithms

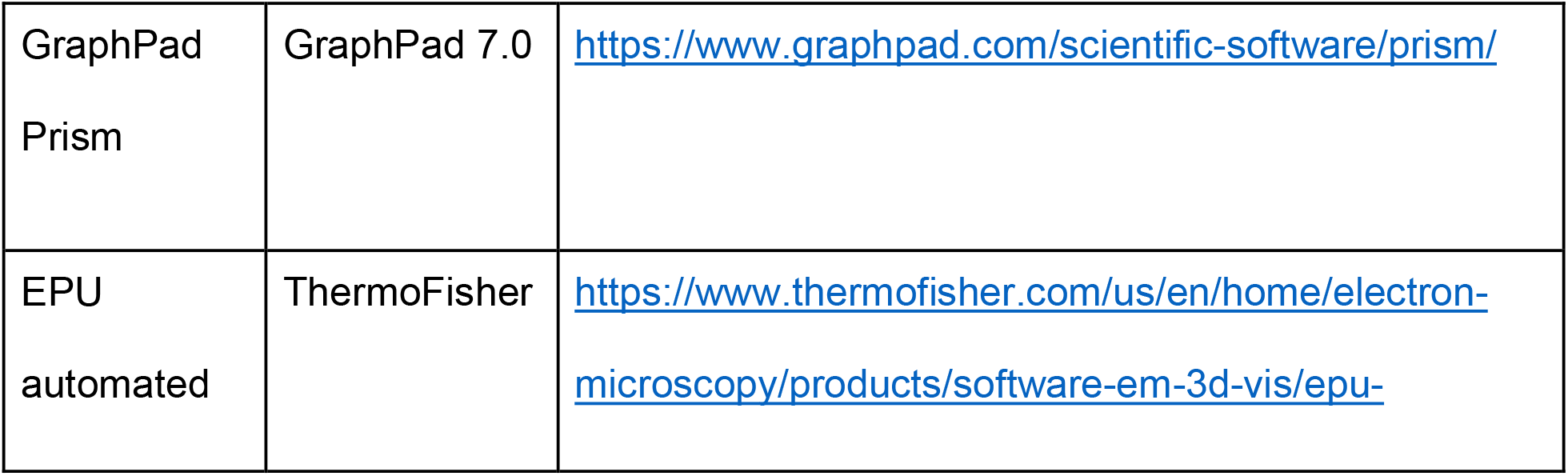
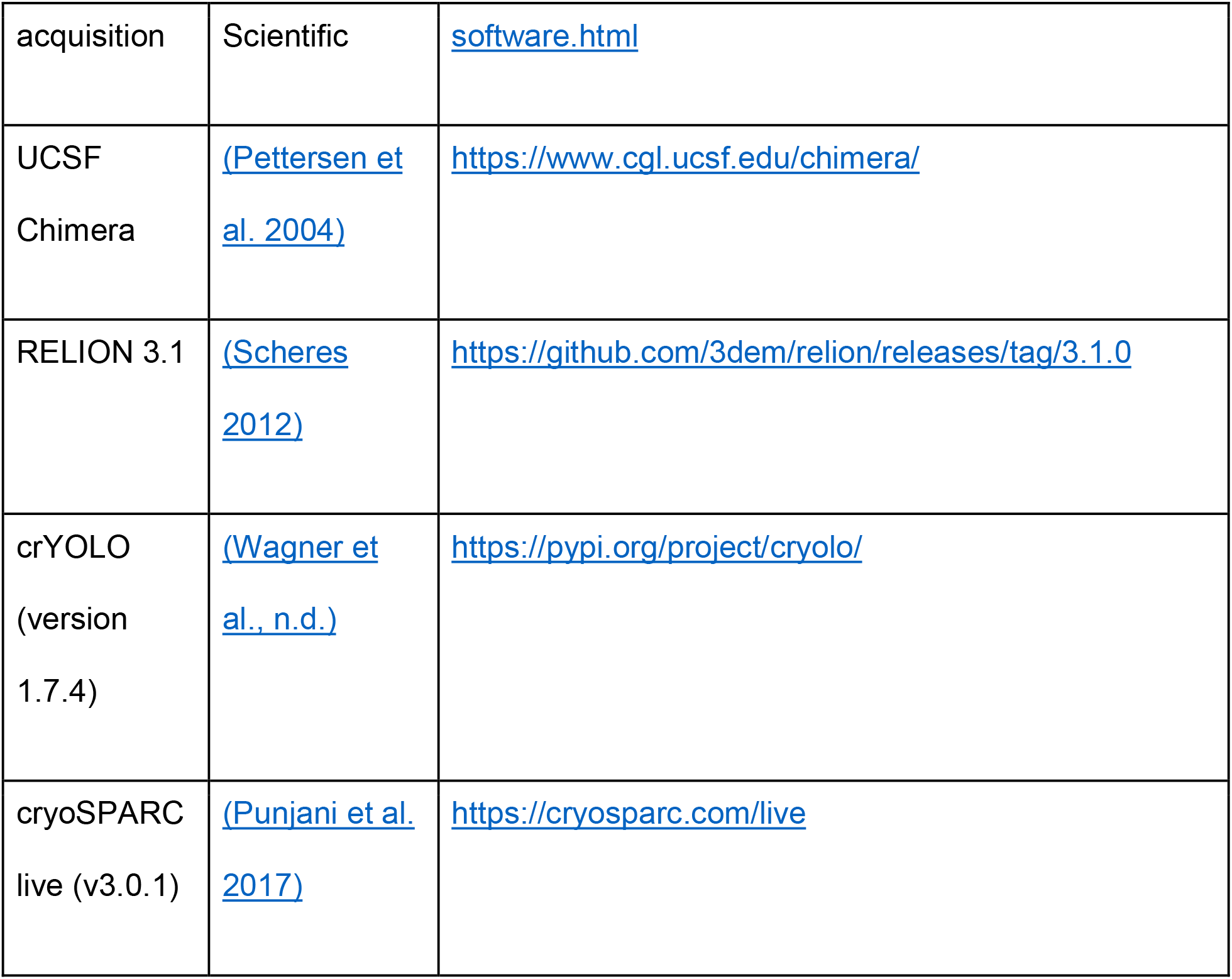

### Method details

#### Cloning, Expression and Purification of Recombinant Spike Protein Constructs

The wild type SARS-CoV-2 S HexaPro expression plasmid was previously described (Hsieh et al. 2020) and was a gift from Jason McLellan (Addgene plasmid #154754; http://n2t.net/addgene:154754; RRID:Addgene_154754).

The VoC RBD mutations were introduced by site-directed mutagenesis (Q5 Site-Directed Mutagenesis Kit, New England Biolabs). Successful cloning was confirmed by Sanger sequencing (Genewiz, Inc.).

Expi293F cells (ThermoFisher, Cat# A14527) were grown in suspension culture using Expi293 Expression Medium (ThermoFisher, Cat# A1435102) at 37°C, 8% CO2. Cells were transiently transfected at a density of 3 x 10^6 cells/mL using linear polyethylenimine (Polysciences Cat# 23966-1). 24-hours following transfection, media was supplemented with 2.2 mM valproic acid and expression carried out for 3-5 days at 37°C, 8% CO2. The supernatant was harvested by centrifugation and filtered through a 0.22 μM filter prior to loading onto a 5 mL HisTrap excel column (Cytiva). The column was washed for 20 CVs with wash buffer (20 mM Tris pH 8.0, 500 mM NaCl), 5 CVs of wash buffer supplemented with 20 mM imidazole and the protein eluted with elution buffer (20 mM Tris pH 8.0, 500 mM NaCl, 500 mM imidazole). Elution fractions containing the protein were pooled and concentrated (Amicon Ultra 100 kDa cut off, Millipore Sigma) for gel filtration. Gel filtration was conducted using a Superose 6 10/300 GL column (Cytiva) pre-equilibrated with GF buffer (20 mM Tris pH 8.0, 150 mM NaCl). Peak fractions corresponding to soluble protein were pooled and concentrated to 4.5 - 5.5 mg/mL (Amicon Ultra 100 kDa cut off, Millipore Sigma). Protein samples were flash-frozen in liquid nitrogen and stored at −80°C.

#### Antibody Production

VH-FC ab8, IgG ab1 and IgG CR3022 were produced as previously described ((Li et al. 2020a) (Li et al. 2020b)). Plasmids encoding light and heavy chains for Fab S309 and S2M11 were synthesized (Synbio). Heavy chains were designed to incorporate a C terminal 6x histidine tag. Expi293 cells were transfected at a density of 3 x 10^6 cells/mL using linear polyethylenimine (Polysciences Cat# 23966-1). 24-hours following transfection, media was supplemented with 2.2 mM valproic acid and expression carried out for 3-5 days at 37°C, 8% CO2. The supernatant was harvested by centrifugation and filtered through a 0.22 μM filter prior to loading onto a 5 mL HisTrap excel column (Cytiva). The column was washed for 20 CVs with wash buffer (20 mM Tris pH 8.0, 500 mM NaCl), 5 CVs of wash buffer supplemented with 20 mM imidazole and the protein eluted with elution buffer (20 mM Tris pH 8.0, 500 mM NaCl, 500 mM imidazole). Elution fractions containing the protein were pooled and concentrated (Amicon Ultra 10 kDa cut off, Millipore Sigma) for gel filtration. Gel filtration was conducted using a Superose 6 10/300 GL column (Cytiva) pre-equilibrated with GF buffer (20 mM Tris pH 8.0, 150 mM NaCl). Peak fractions corresponding to soluble protein were pooled and concentrated to 8 - 20 mg/mL (Amicon Ultra 10 kDa cut off, Millipore Sigma). Protein samples were stored at 4°C until use.

#### Electron Microscopy Sample Preparation and Data Collection

S-protein samples were prepared at 2.25 mg/mL, with and without the addition of ACE2 (~1:1.25 S-protein trimer:ACE2 molar ratio) (New England Biolabs). Vitrified samples of S-protein constructs with and without ACE2 were prepared by first glow discharging Quantifoil R1.2/1.3 300 mesh holey carbon copper grids for 1 minute using a Pelco easiGlow glow discharge unit (Ted Pella) and then applying 1.8 μL of protein suspension to the surface of the grid. Grids were blotted (12 sec, blot force −10) and plunge frozen into liquid ethane using a Vitrobot Mark IV (Thermo Fisher Scientific) at a temperature of 10 °C and a humidity level of 100%. All cryo-EM samples were imaged using a 300 kV Titan Krios G4 transmission electron microscope (ThermoFisher Scientific) equipped with a Falcon4 direct electron detector in electron event registration (EER) mode. Movies were collected at 155,000x magnification (physical pixel size 0.5 Å) over a defocus range of −0.5 μm to −3 μm with a total dose of 40 e-/ Å^2^ using EPU automated acquisition software.

#### Image Processing

In general, all data processing was performed in cryoSPARC v.3.0.1 (Punjani et al., 2017) unless stated otherwise. Motion correction in patch mode (EER upsampling factor 1, EER number of fractions 40), CTF estimation in patch mode, reference-free particle picking and particle extraction were performed on-the-fly in cryoSPARC. After preprocessing, particles were subjected to 2D classification and/or 3D heterogeneous classification. Final 3D refinement was done with per particle CTF estimation and aberration correction. For complex of spike protein ectodomain and human ACE2, focused refinements were performed with a soft mask covering single RBD and its bound ACE2. Global resolution and focused resolution were according to the gold-standard FSC (Bell et al., 2016).

#### Model Building and Refinement

For models of spike protein ectodomain alone, SARS-CoV-2 HexaPro S trimer with N501Y mutation (PDB code 7MJG) were docked into cryo-EM density using UCSF Chimera v.1.15 (Pettersen et al., 2004). Then mutation and manual adjustment were done with COOT v.0.9.3 (Emsley et al., 2010), followed by iterative rounds of refinement in COOT and Phenix v.1.19 (Afonine et al., 2018). Glycans were added at N-linked glycosylation sites in COOT. For models of complex of spike protein ectodomain and human ACE2, the RBD-ACE2 subcomplex was built using coordinates of PDB code 7MJN as initial model and refined against focused refinement maps. Then it was docked into global refinement maps together with individual domains of spike protein. Model validation was performed using MolProbity (Chen et al., 2010). Figures were prepared using UCSF Chimera, UCSF ChimeraX v.1.1.1 (Goddard et al., 2018), and PyMOL (v.2.2 Schrodinger, LLC).

#### Biolayer Interferometry (BLI) S protein-ACE2 Binding Assay

The binding kinetics of SARS-CoV-2 trimers and human ACE2 was analyzed with the biolayer interferometer BLItz (ForteBio, Menlo Park, CA). Protein-A biosensors (ForteBio: 18–5010) were coated with ACE2-mFc (40 μg/mL) for 2 min and incubated in DPBS (pH = 7.4) to establish baselines. Concentrations of 125, 250, 500 and 1000 nM spike trimers were used for association for 2 min followed by dissociation in DPBS for 5 min. The association (k_on_) and dissociation (k_off_) rates were derived from the sensorgrams fitting and used to calculate the binding equilibrium constant (K_D_).

#### Enzyme-Linked Immunosorbent Assay (ELISA)

100 μl of wild-type or VoC RBD mutant SARS-CoV-2 S protein preparations were coated onto 96-well MaxiSorp^™^ plates at 2 μg/ml in PBS overnight at 4°C. All washing steps were performed 5 times with PBS + 0.05% Tween 20 (PBS-T). After washing, wells were either incubated with blocking buffer (PBS-T + 2% BSA) for 1 hr at room temperature. After washing, wells were incubated with dilutions of primary antibodies in PBS-T + 0.5% BSA buffer for 1 hr at room temperature. After washing, wells were incubated with goat anti-human IgG (Jackson ImmunoResearch) at a 1:8,000 dilution in PBS-T + 0.5% BSA buffer for 1 hr at room temperature. After washing, the substrate solution (Pierce^™^ 1-Step^™^) was used for colour development according to the manufacturer’s specifications. Optical density at 450 nm was read on a Varioskan Lux plate reader (Thermo Fisher Scientific).

#### Analysis of Convalescent Patient Antibody Footprints

PDB entries of SARS-CoV-2 spike or RBD complexes with antibody fragments isolated from convalescent patients were selected. Antibody footprints were determined by consulting respective depositing studies along with analysis of protein-protein contacts using PDBsum (Laskowski et al., 2018).

