## Supplemental Figures for "Structural Analysis of Receptor Binding Domain Mutations in SARS-CoV-2 Variants of Concern that Modulate ACE2 and Antibody Binding"

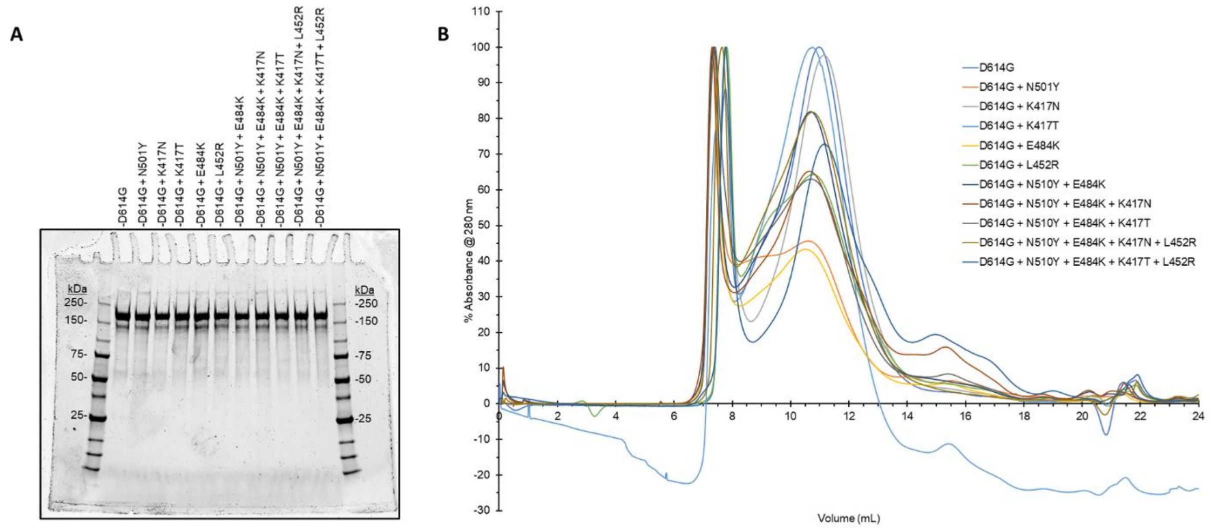


**Figure S1. Purification of SARS-CoV-2 S-proteins containing VoC RBD mutations. (A)** SDS Polyacrylamide gel electrophoresis (SDS-PAGE) gel of the eleven SARS-CoV-2 RBD mutant S-proteins employed in this study. Normalized amounts (2 µg) of each variant spike was loaded in each well. **(B)** Superose 6 10/300 GL size-exclusion traces of the SARS-CoV-2 RBD mutant S-proteins.

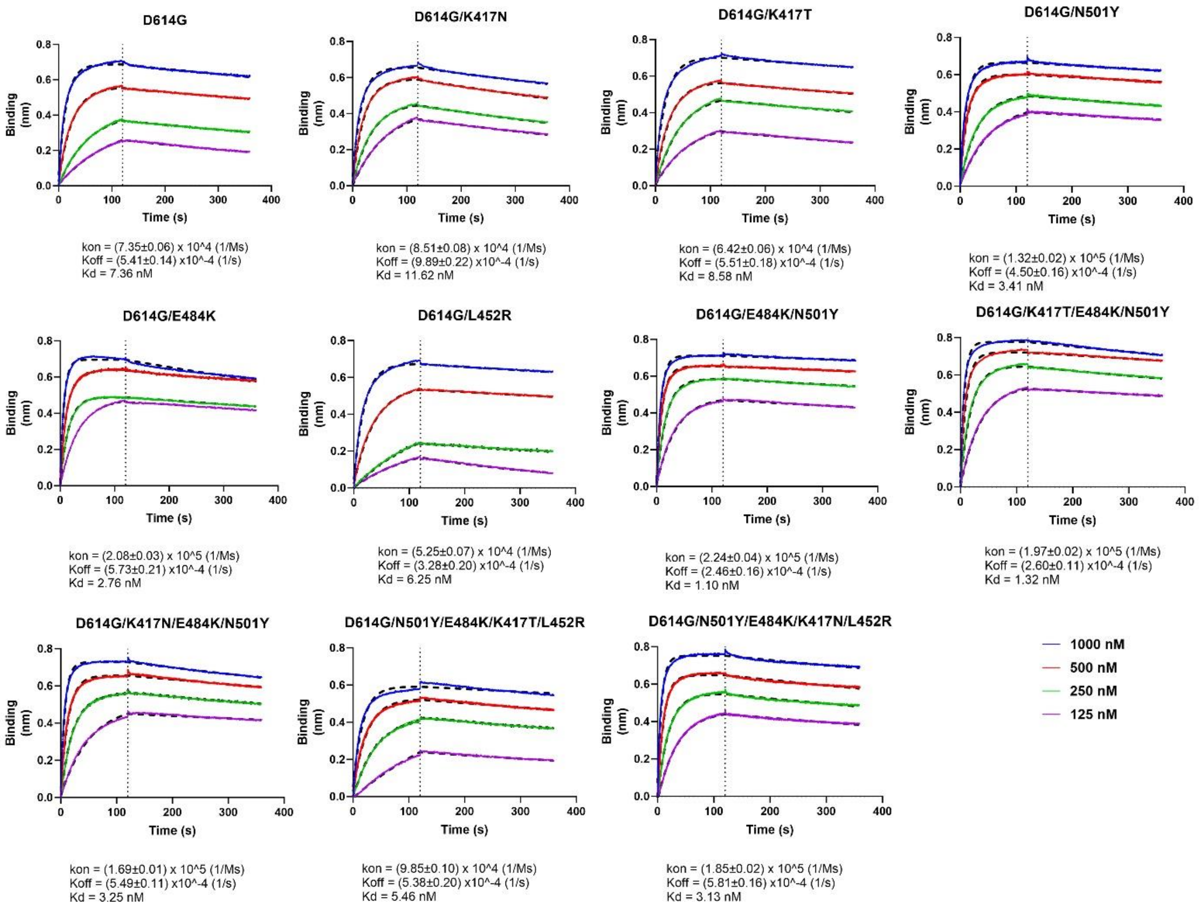


**Figure S2. Raw Biolayer interferometry (BLI) sensorgrams for all spike constructs included in this study, related to figure 2.** Wild-type (D614G) and variant spikes were assessed for binding of immobilized ACE2 at increasing concentrations as indicated. Shown is the extent of binding as determined by shift in wavelength (nm: nanometers). Biophysical parameters ($K_D$, $k_{on}$, $k_{off}$) are shown as means. Errors correspond to standard deviations.

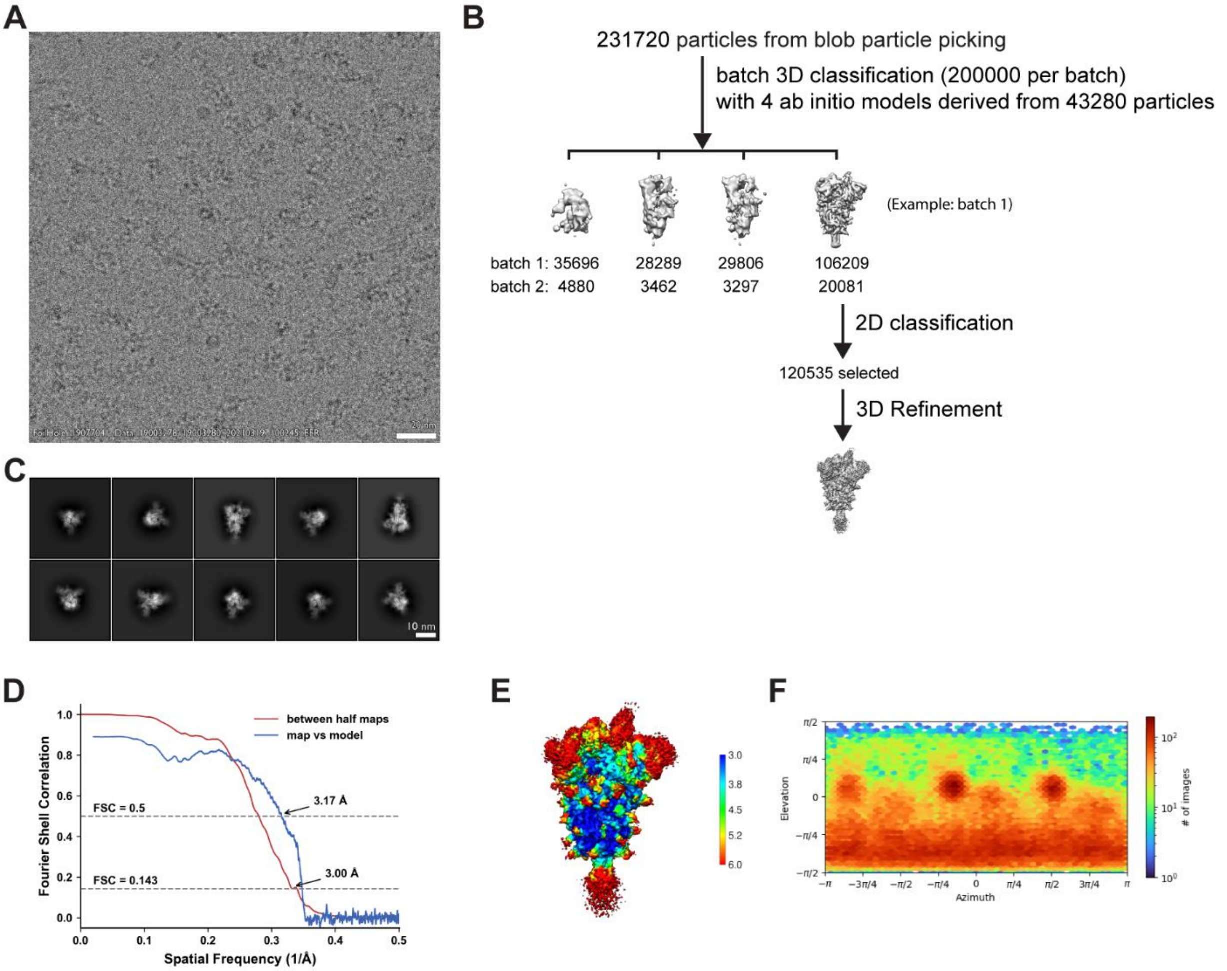


**Figure S3. Cryo-EM data processing and validation for the D614G spike protein ectodomain. (A)** Representative cryo-EM micrograph. **(B)** Workflow of cryo-EM image processing. **(C)** Representative 2D classes. **(D)** FSC curves. **(E)** Local resolution. **(F)** Viewing direction distribution plot.

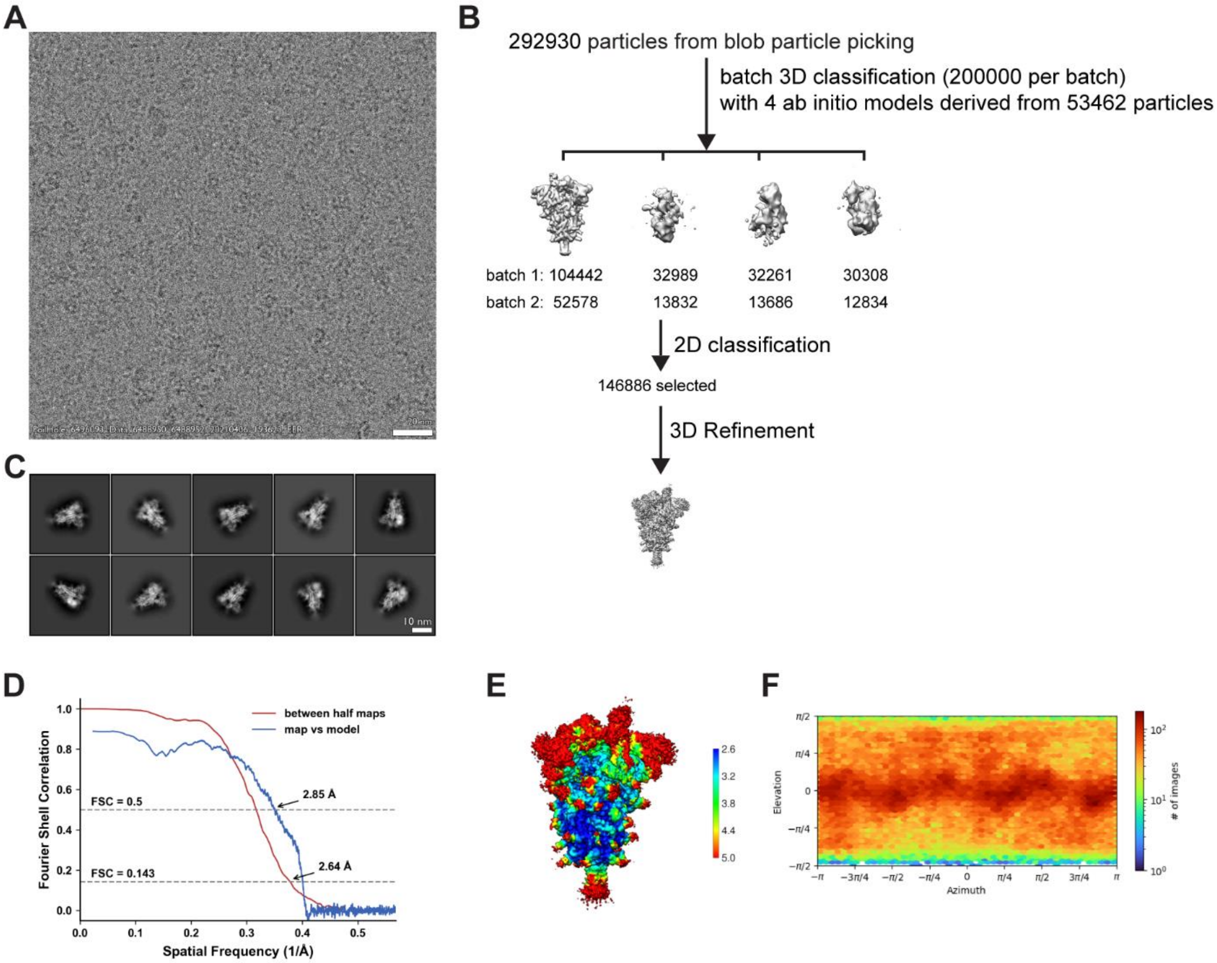


**Figure S4. Cryo-EM data processing and validation for the D614G + L452R spike protein ectodomain.** **(A)** Representative cryo-EM micrograph. **(B)** Workflow of cryo-EM image processing. **(C)** Representative 2D classes. **(D)** FSC curves. **(E)** Local resolution. **(F)** Viewing direction distribution plot.

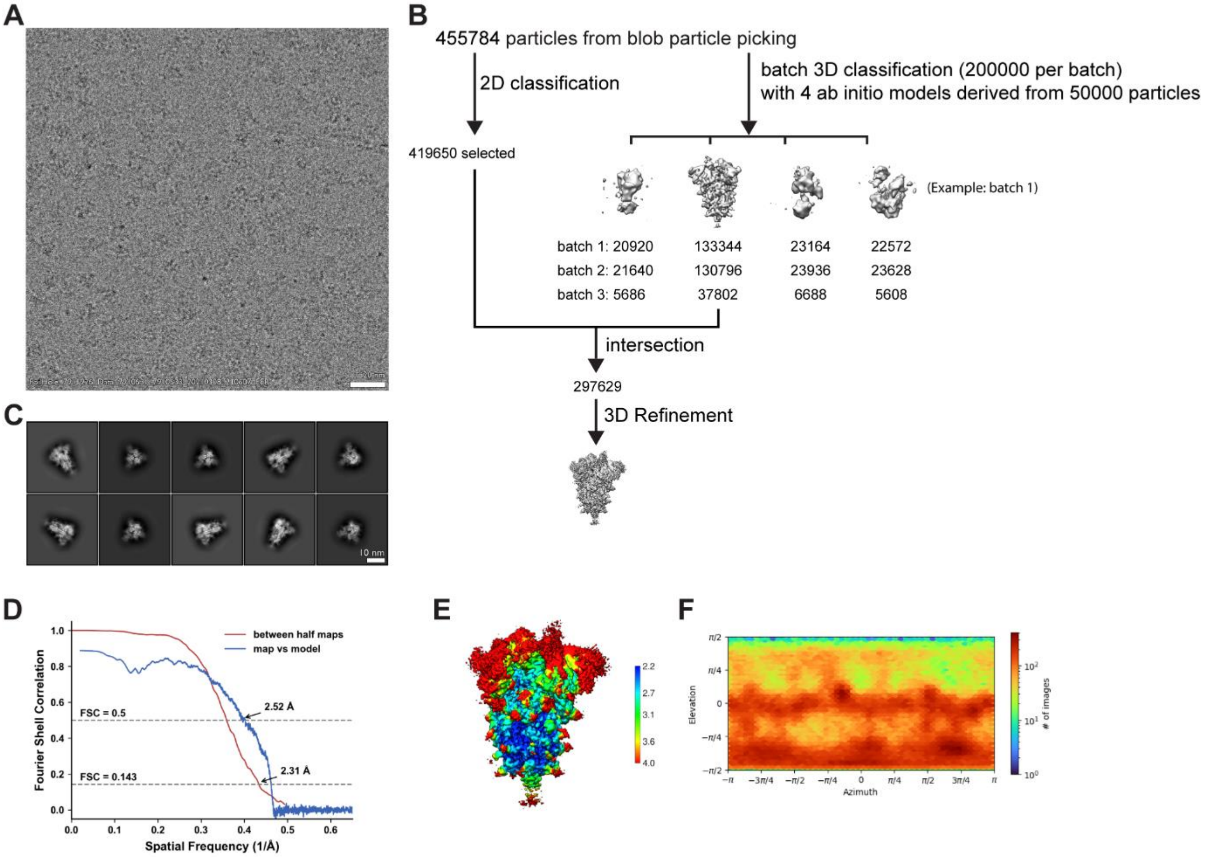


**Figure S5. Cryo-EM data processing and validation for the D614G + N501Y spike protein ectodomain. (A)** Representative cryo-EM micrograph. **(B)** Workflow of cryo-EM image processing. **(C)** Representative 2D classes. **(D)** FSC curves. **(E)** Local resolution. **(F)** Viewing direction distribution plot

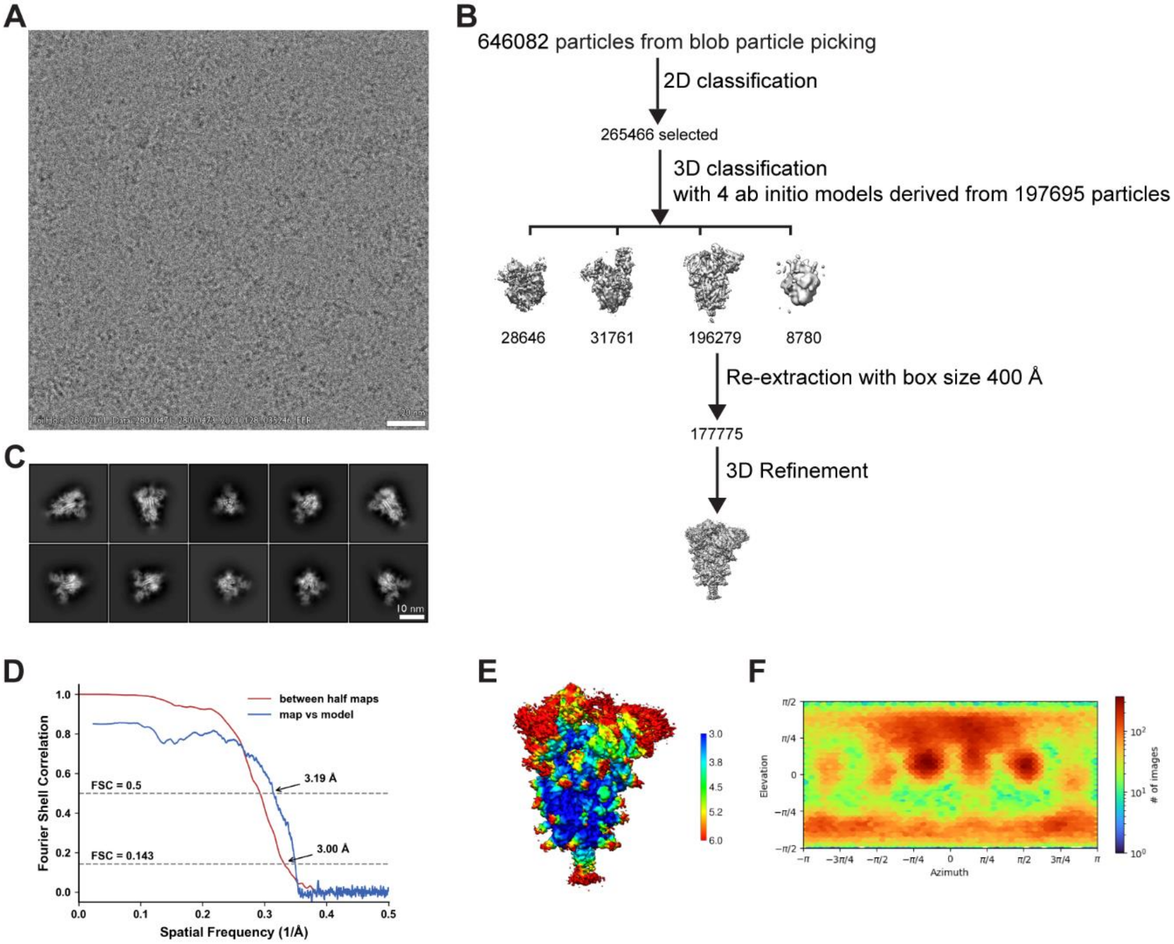


**Figure S6. Cryo-EM data processing and validation for the D614G + N501Y + E484K spike protein ectodomain. (A)** Representative cryo-EM micrograph. **(B)** Workflow of cryo-EM image processing. **(C)** Representative 2D classes. **(D)** FSC curves. **(E)** Local resolution. **(F)** Viewing direction distribution plot

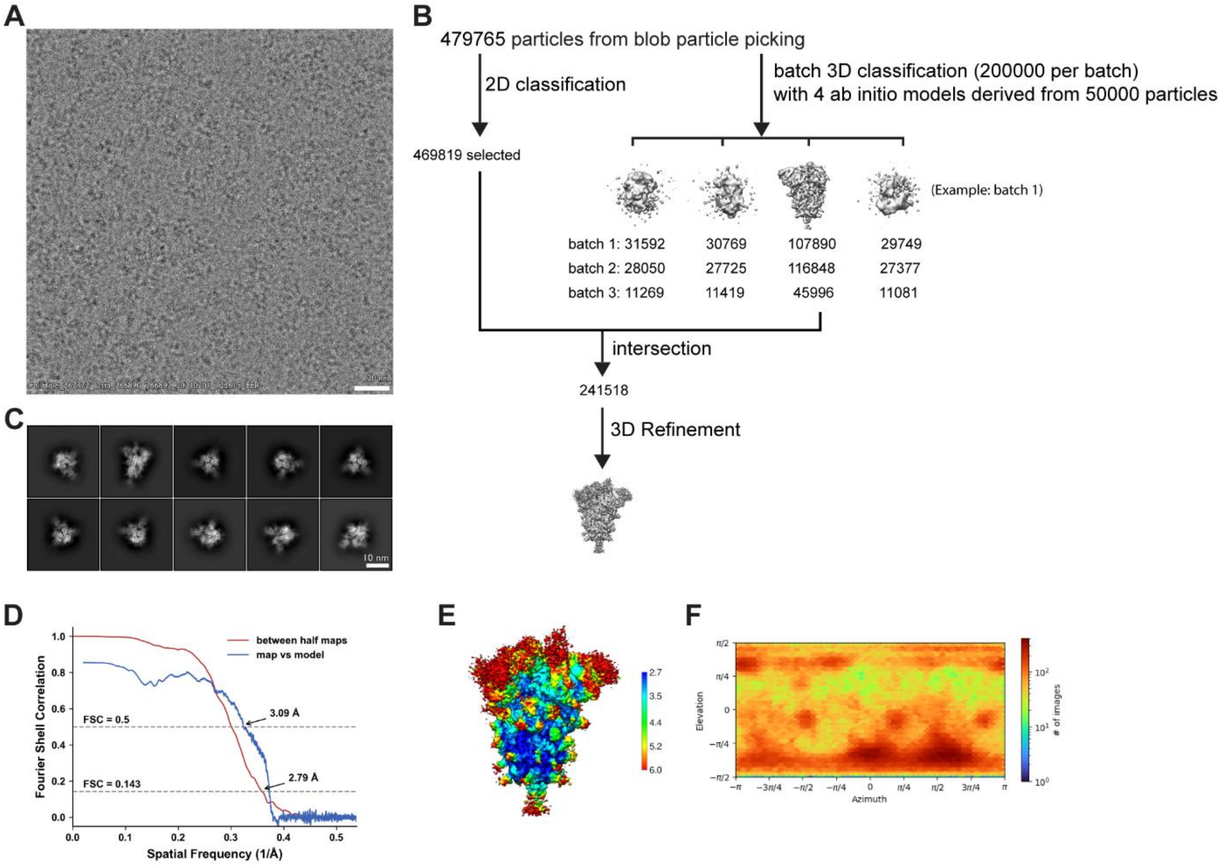


**Figure S7. Cryo-EM data processing and validation for the D614G + N501Y + E484K + K417N spike protein ectodomain. (A)** Representative cryo-EM micrograph. **(B)** Workflow of cryo-EM image processing. **(C)** Representative 2D classes. **(D)** FSC curves. **(E)** Local resolution. **(F)** Viewing direction distribution plot

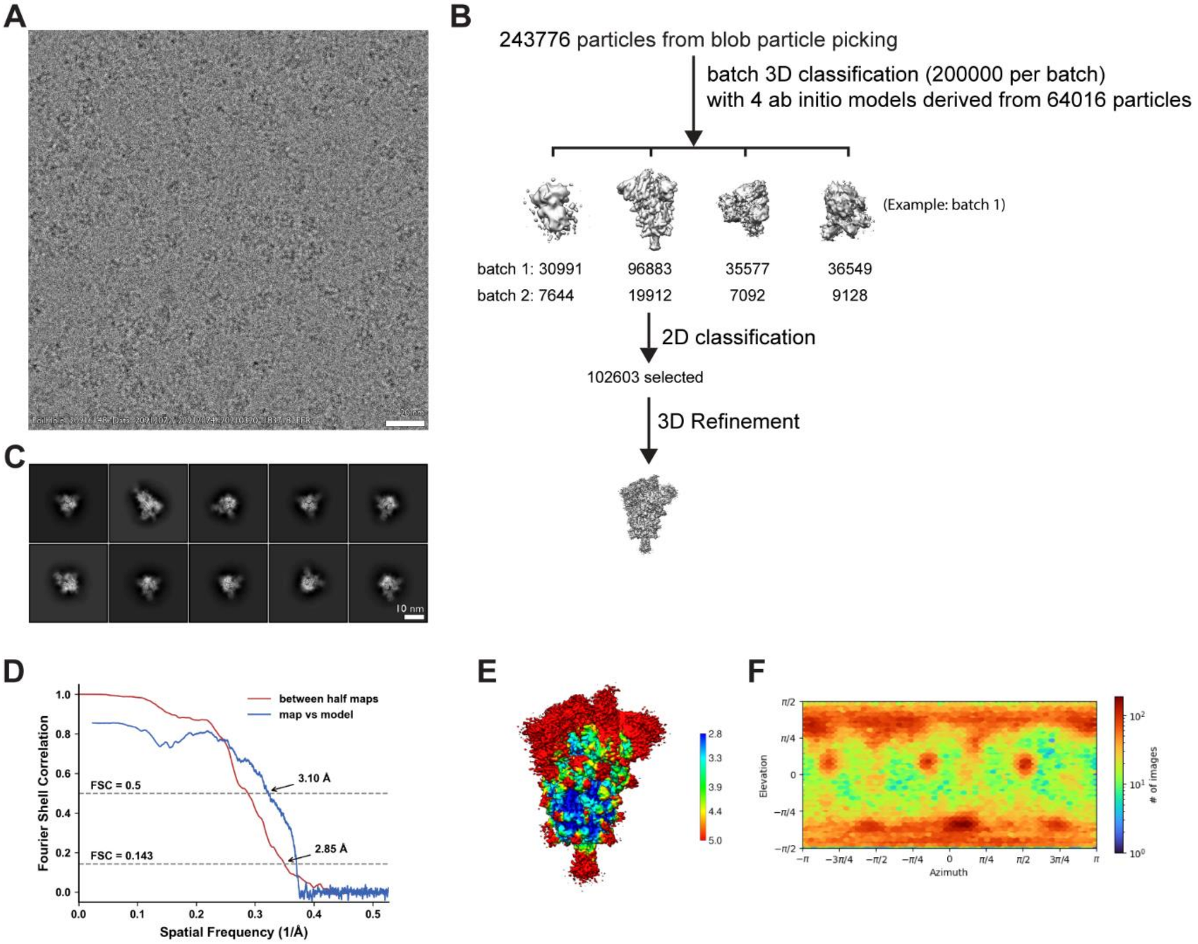


**Figure S8. Cryo-EM data processing and validation for the D614G + N501Y + E484K + K417T spike protein ectodomain. (A)** Representative cryo-EM micrograph. **(B)** Workflow of cryo-EM image processing. **(C)** Representative 2D classes. **(D)** FSC curves. **(E)** Local resolution. **(F)** Viewing direction distribution plot

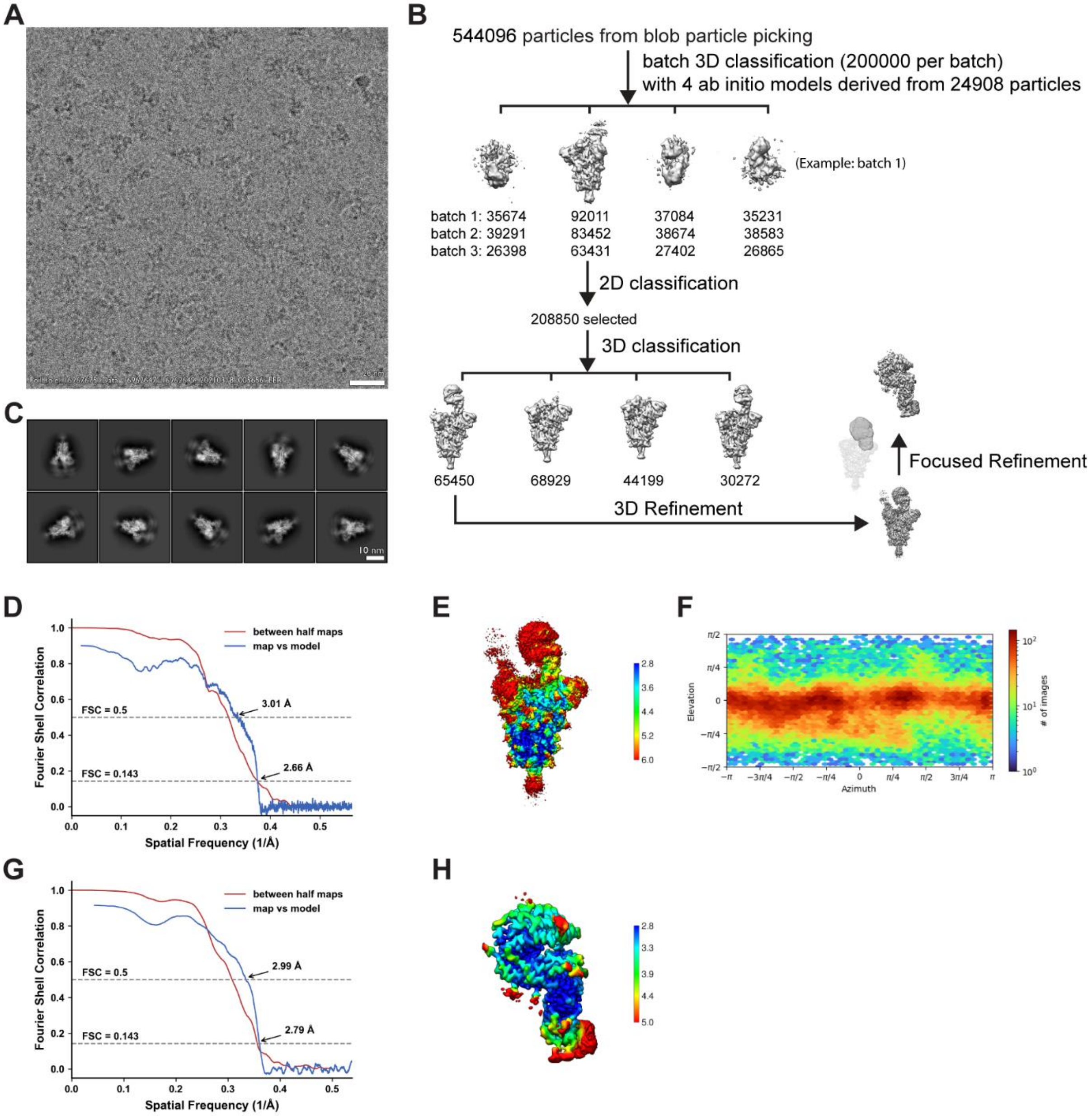


**Figure S9. Cryo-EM data processing and validation for complex of D614G spike protein ectodomain and human ACE2. (A)** Representative cryo-EM micrograph. **(B)** Workflow of cryo-EM image processing. **(C)** Representative 2D classes. **(D-F)** FSC curves **(D)**, local resolution **(E)** and viewing direction distribution plot **(F)** of global refinement. **(G-H)** FSC curves **(G)** and local resolution **(H)** of focused refinement.

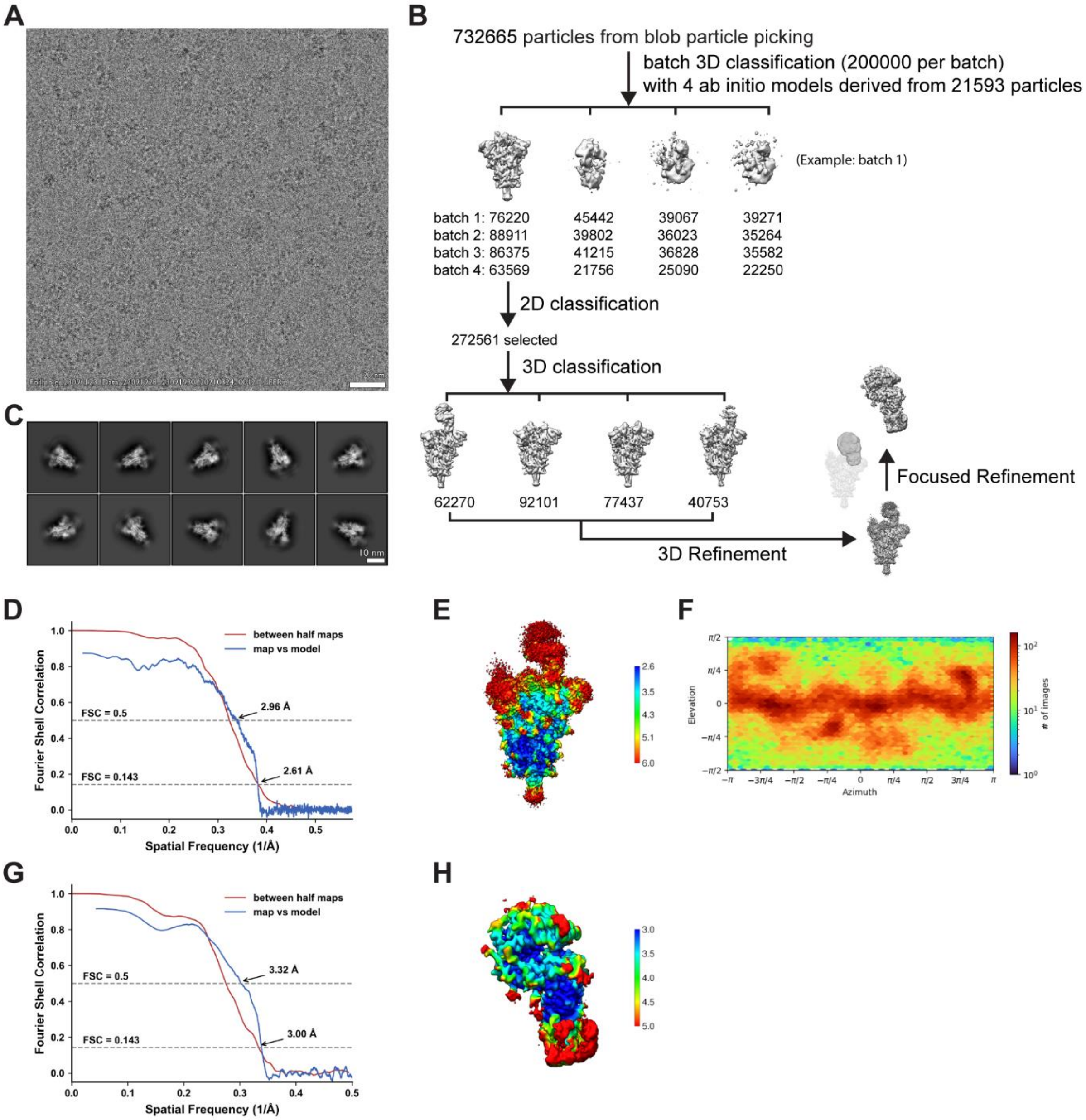


**Figure S10. Cryo-EM data processing and validation for complex of D614G + L452R spike protein ectodomain and human ACE2. (A)** Representative cryo-EM micrograph. **(B)** Workflow of cryo-EM image processing. **(C)** Representative 2D classes. **(D-F)** FSC curves **(D)**, local resolution **(E)** and viewing direction distribution plot **(F)** of global refinement. **(G-H)** FSC curves **(G)** and local resolution **(H)** of focused refinement.

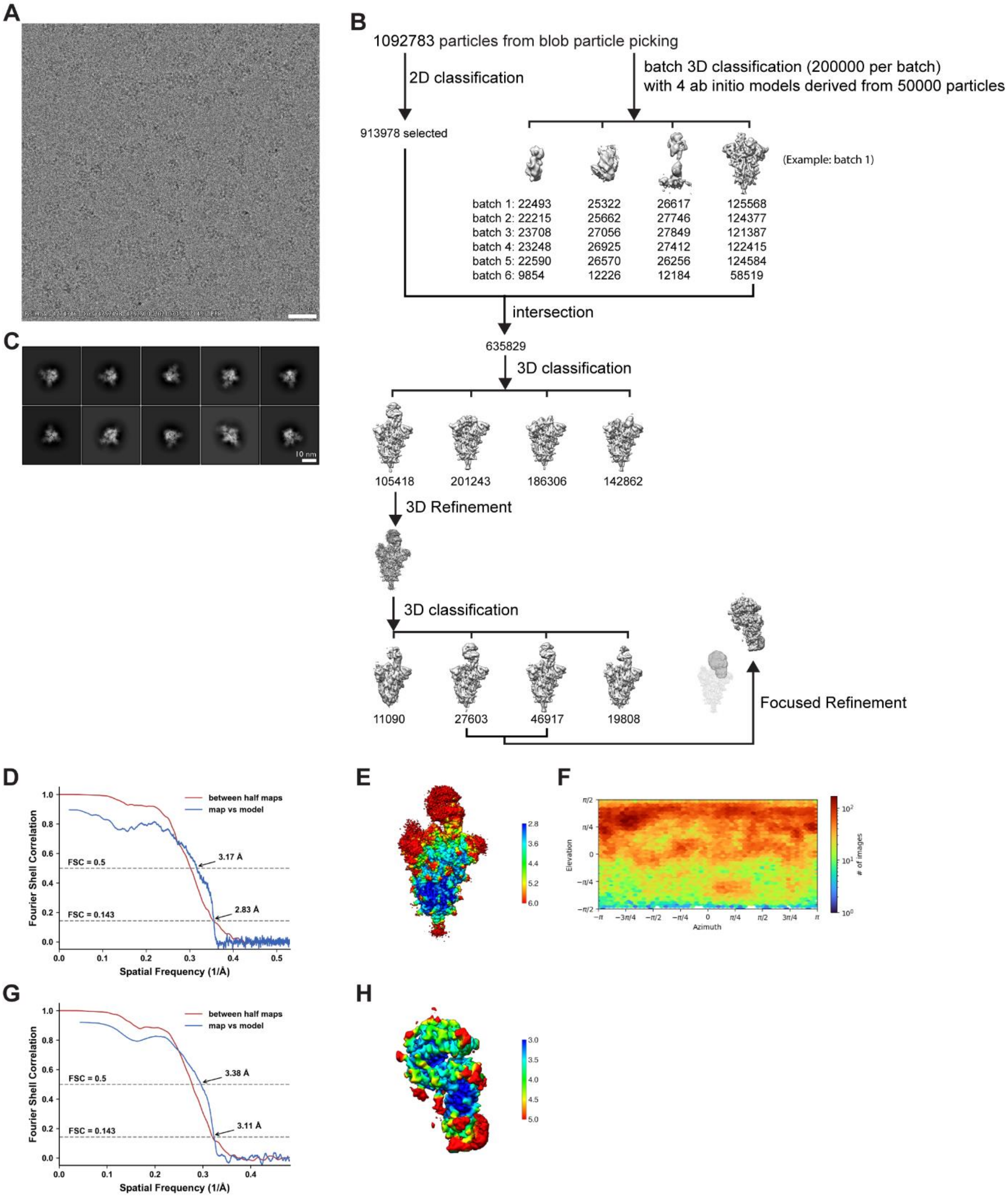


**Figure S11. Cryo-EM data processing and validation for complex of D614G + N501Y spike protein ectodomain and human ACE2. (A)** Representative cryo-EM micrograph. **(B)** Workflow of cryo-EM image processing. **(C)** Representative 2D classes. **(D-F)** FSC curves **(D)**, local resolution **(E)** and viewing direction distribution plot **(F)** of global refinement. **(G-H)** FSC curves **(G)** and local resolution **(H)** of focused refinement.

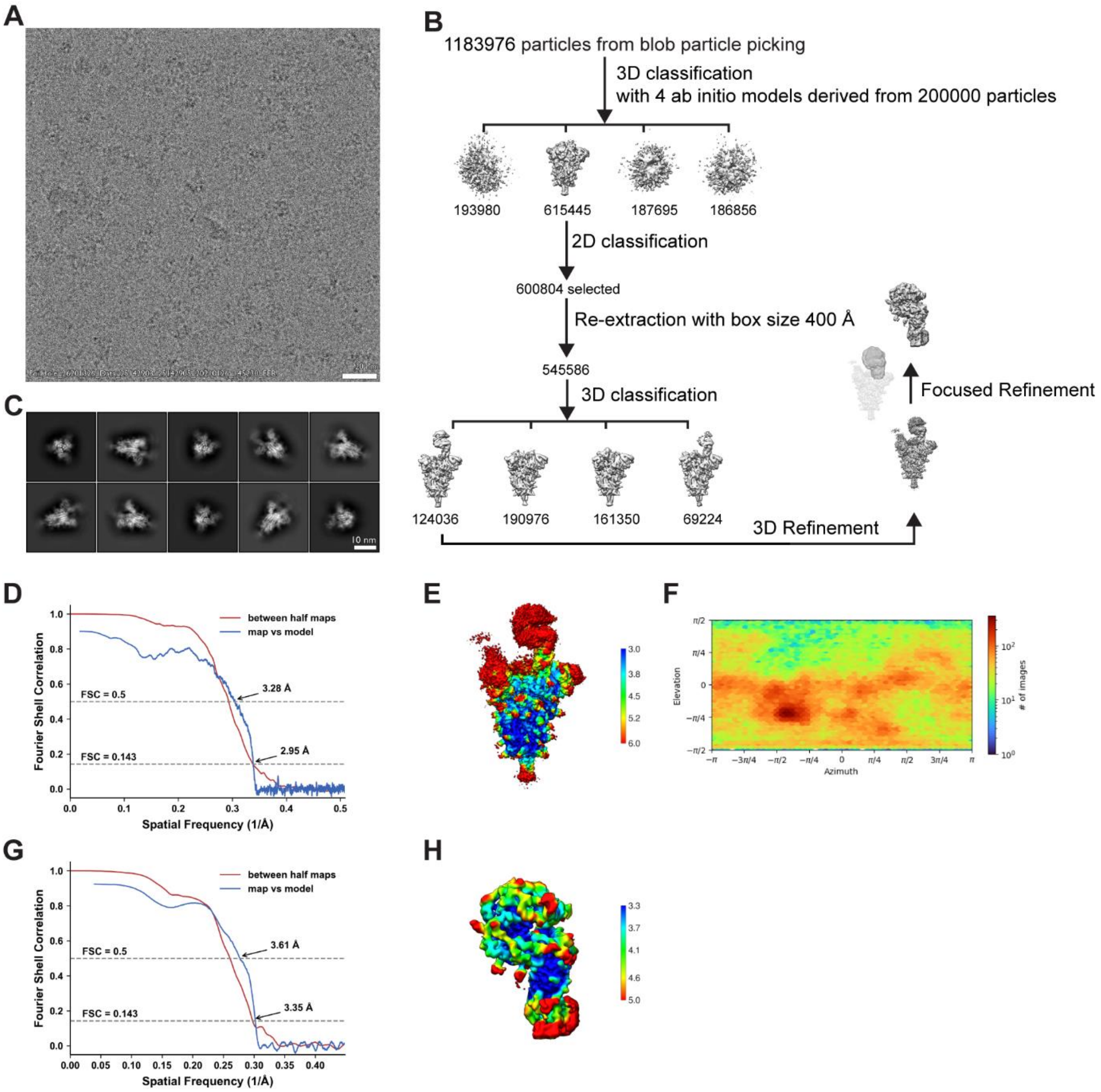


**Figure S12. Cryo-EM data processing and validation for complex of D614G + N501Y + E484K spike protein ectodomain and human ACE2. (A)** Representative cryo-EM micrograph. **(B)** Workflow of cryo-EM image processing. **(C)** Representative 2D classes. **(D-F)** FSC curves **(D)**, local resolution **(E)** and viewing direction distribution plot **(F)** of global refinement. **(G-H)** FSC curves **(G)** and local resolution **(H)** of focused refinement.

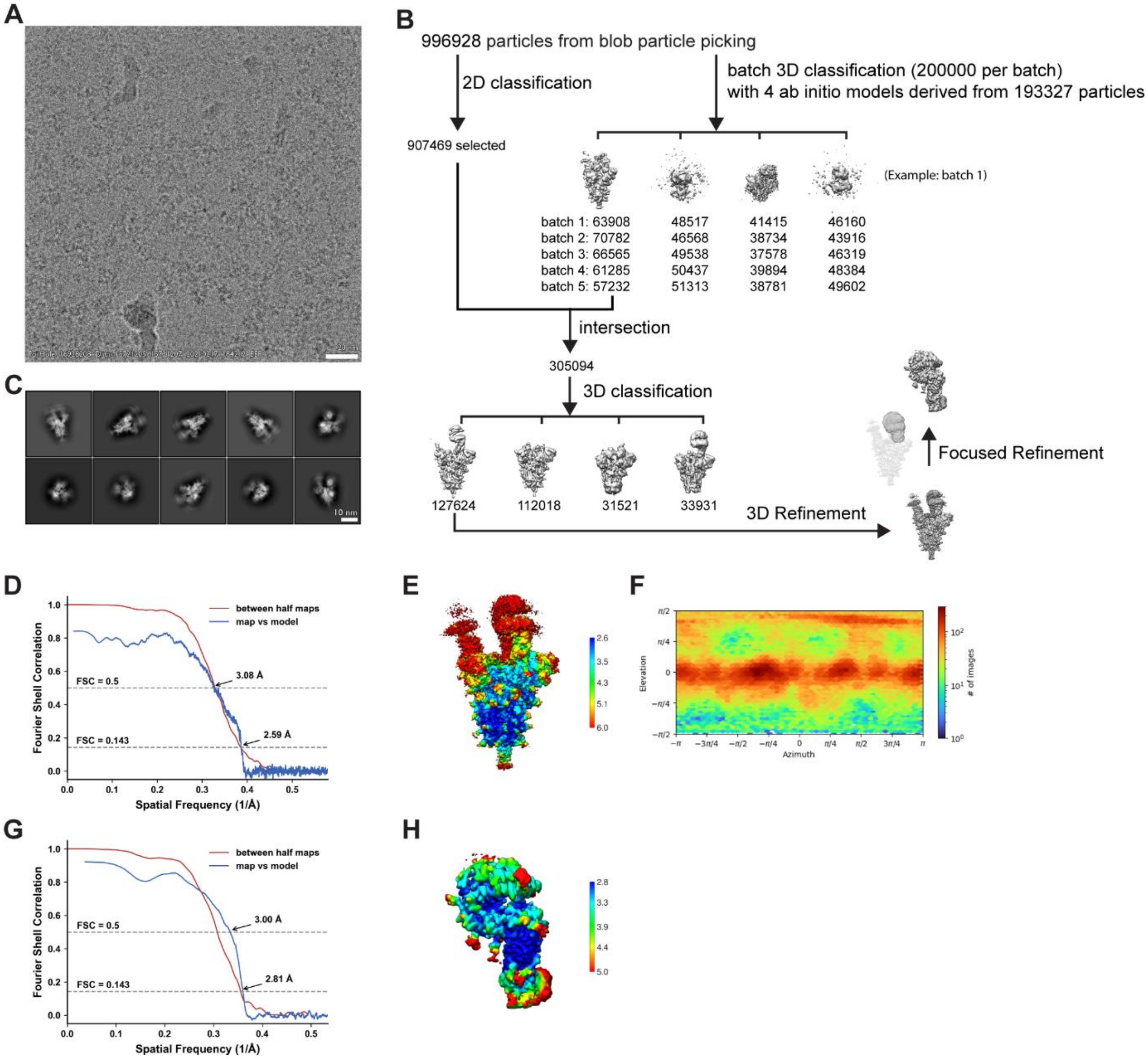


**Figure S13. Cryo-EM data processing and validation for complex of D614G + N501Y + E484K + K417N spike protein ectodomain and human ACE2. (A)** Representative cryo-EM micrograph. **(B)** Workflow of cryo-EM image processing. **(C)** Representative 2D classes. **(D-F)** FSC curves **(D)**, local resolution **(E)** and viewing direction distribution plot **(F)** of global refinement. **(G-H)** FSC curves **(G)** and local resolution **(H)** of focused refinement.

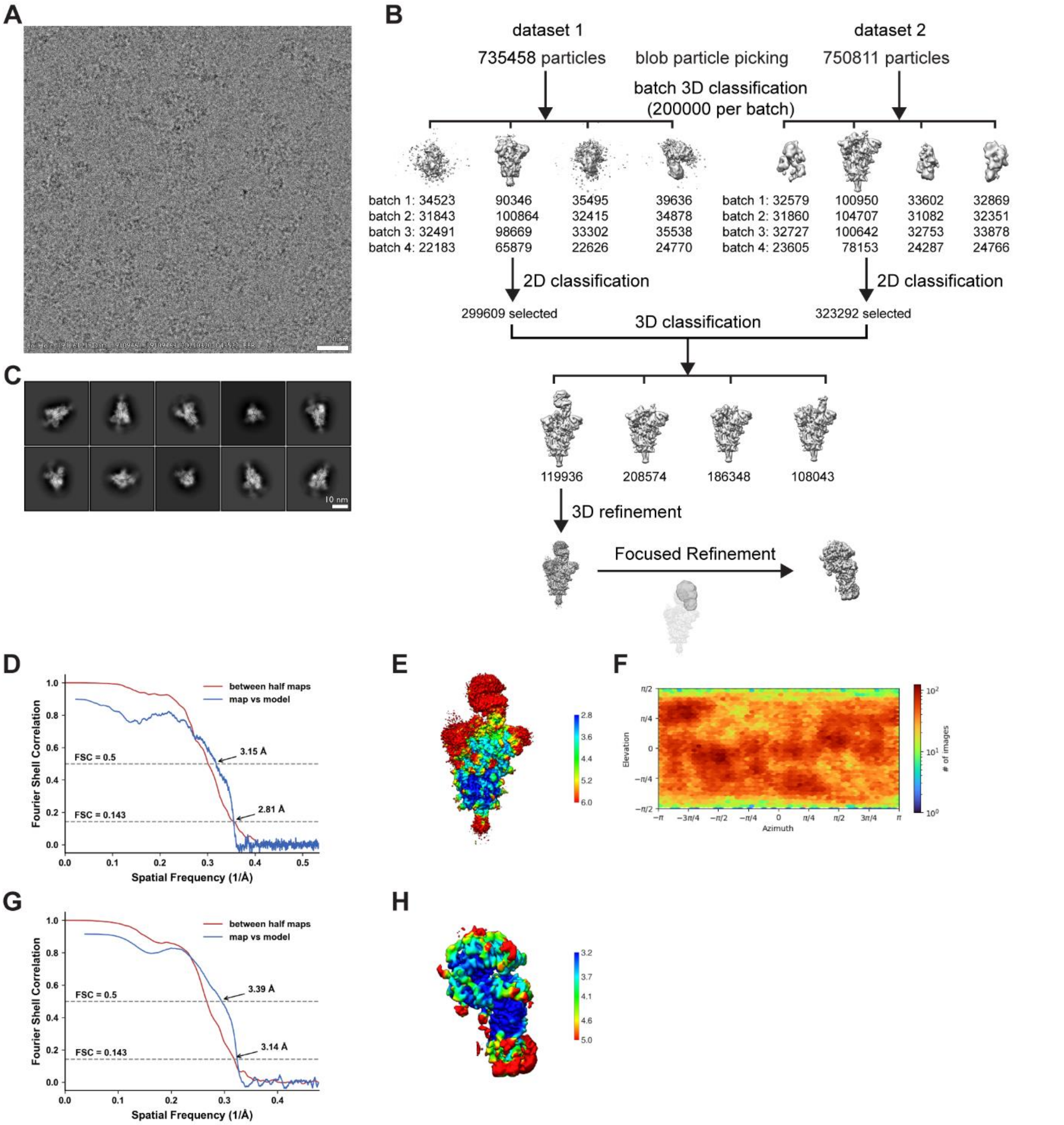


**Figure S14. Cryo-EM data processing and validation for complex of D614G + N501Y + E484K + K417T spike protein ectodomain and human ACE2. (A)** Representative cryo-EM micrograph. **(B)** Workflow of cryo-EM image processing. **(C)** Representative 2D classes. **(D-F)** FSC curves **(D)**, local resolution **(E)** and viewing direction distribution plot **(F)** of global refinement. **(G-H)** FSC curves **(G)** and local resolution **(H)** of focused refinement.

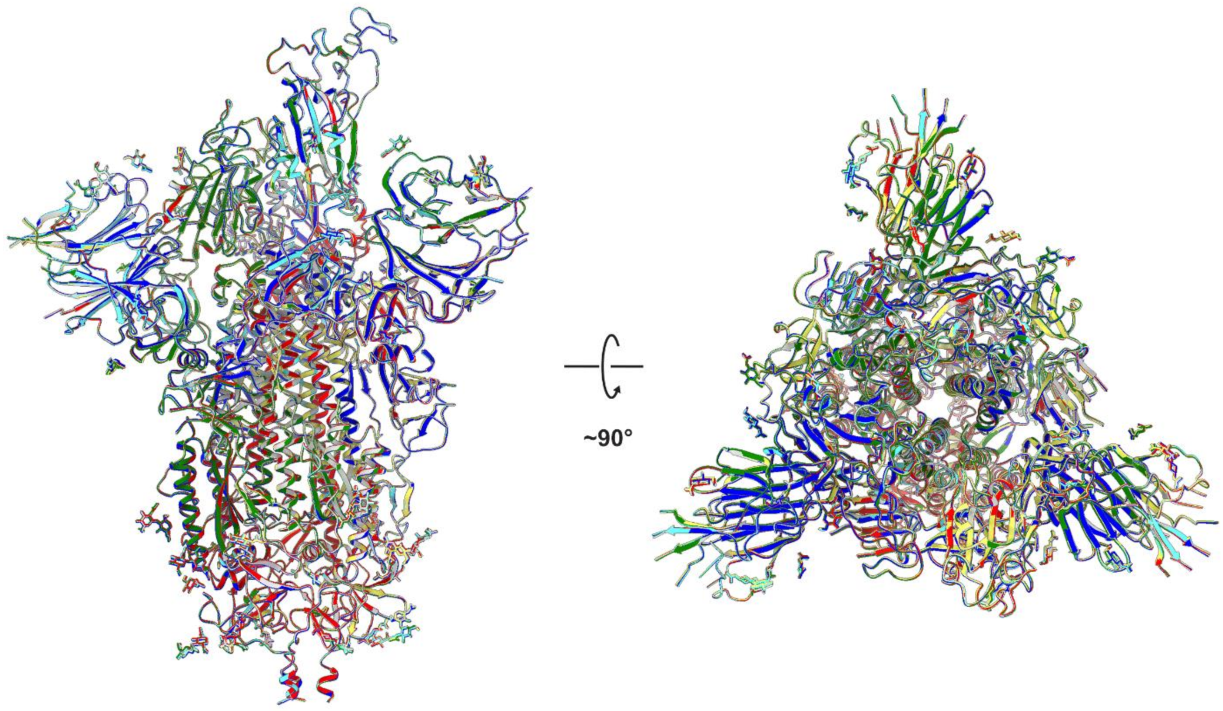


**Figure S15. Superposition of all mutant ectodomain structures characterized in this study.** Gray: D614G, Red: D614G + N501Y, Green: D614G + N501Y + E484K, Blue: D614G + N501Y + E484K + K417N, Yellow: D614G + N501Y + E484K + K417T, Cyan: D614G + L452R

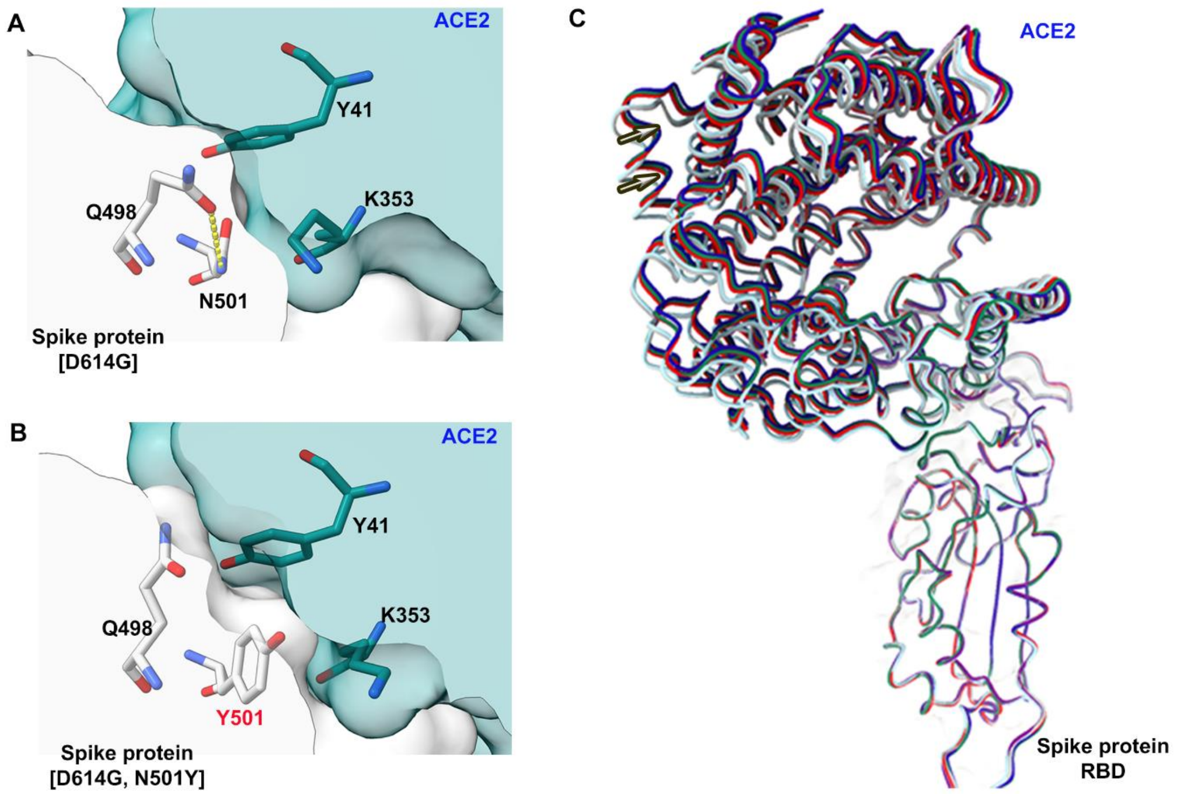


**Figure S16. Structural effects of the N501Y mutation on RBD-ACE2 complexes. (A)** Zoomed in view of position 501 and adjacent residues at the D614G RBD-ACE2 interface. **(B)** Zoomed in view of position 501 and adjacent residues at the D614G, N501Y RBD-ACE2 interface. **(C)** Superposition of the RBD in the RBD-ACE2 structures from all complexes reported in this study. Cyan and grey models correspond to structures harboring N at position 501, all other models correspond to structures harboring Y and position 501. Movement of the ACE2 helix distal to the RBD binding interface is highlighted with arrows.

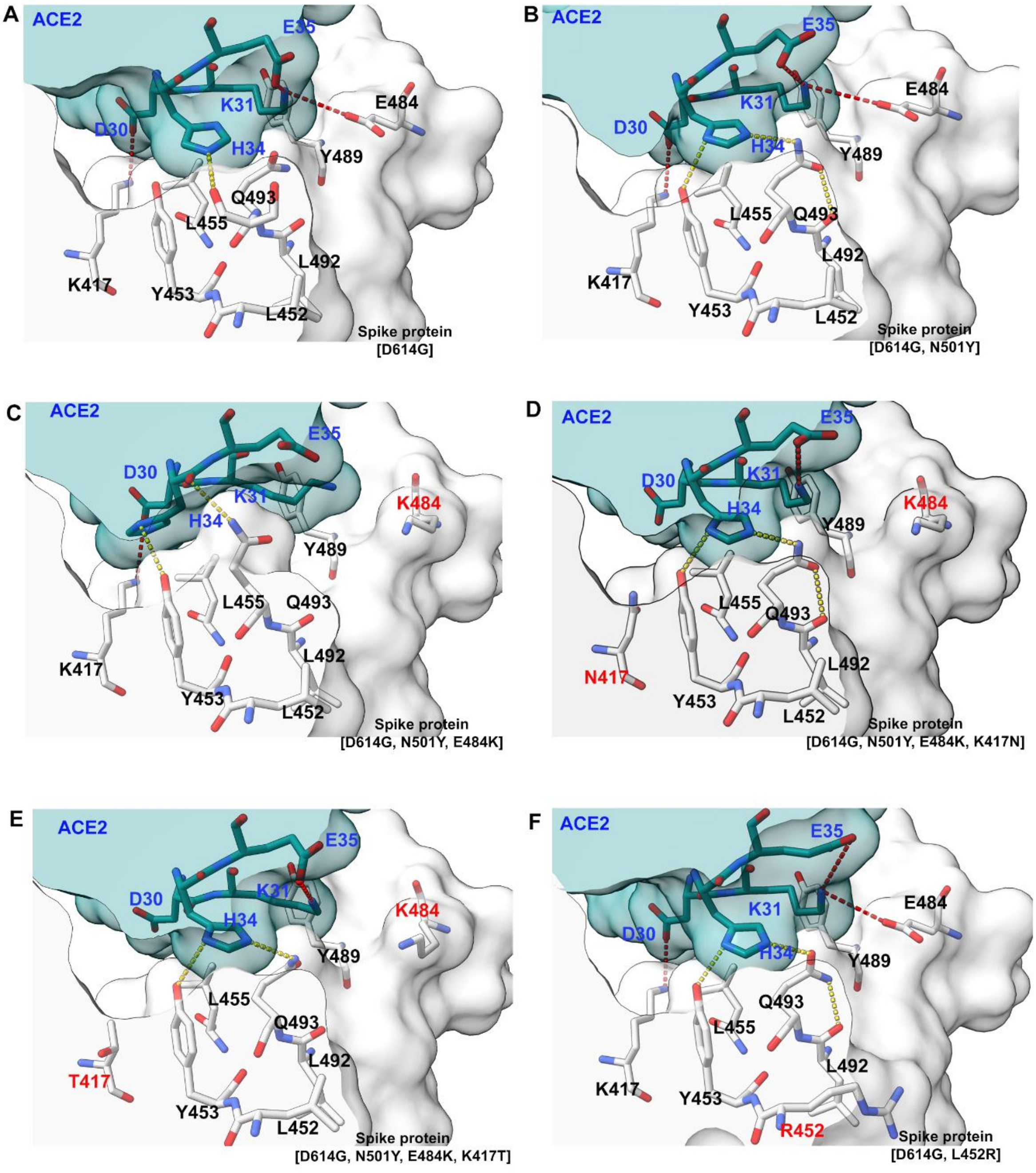


**Figure S17. Analysis of RBD-ACE2 interactions. (A – F)** Zoomed in views of residue H34 within ACE2, residue Q493 within the RBD, and adjacent residues at the RBD-ACE2 interface for all complexes studied. Mutated residues are highlighted in red. Hydrogen bonds are shown as dotted yellow lines, electrostatic interactions are shown as red dotted lines.

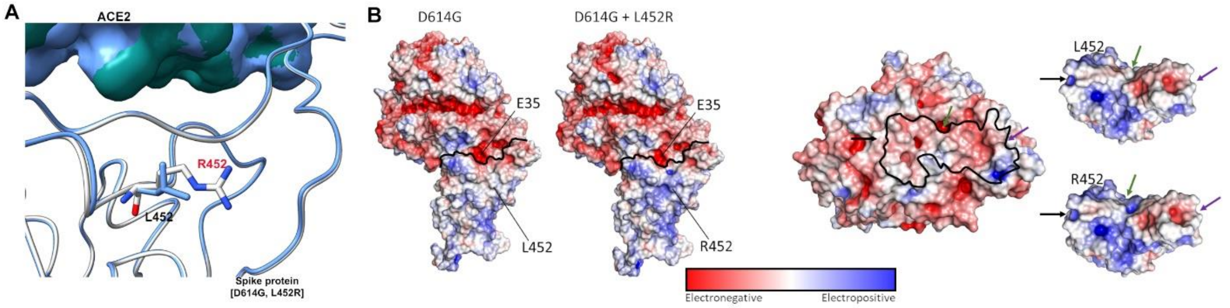


**Figure S18. L452R enhances electrostatic complementarity between ACE2 and the SARS-CoV-2 RBD.** **(A)** Superposition of D614G (blue) and D614G +L452R (white) RBD-ACE2 interfaces. **(B)** Left: Electrostatic surface representations of CryoEM structures of wild-type (D614G) and L452R RBD-ACE2 complexes. Right: Electrostatic surface representation of ACE2, wild-type (D614G) RBD and L452R RBD in isolation. The RBD binding interface on ACE2 is outlined in black and arrows for orientation are provided for both ACE2 and RBD surface representations.

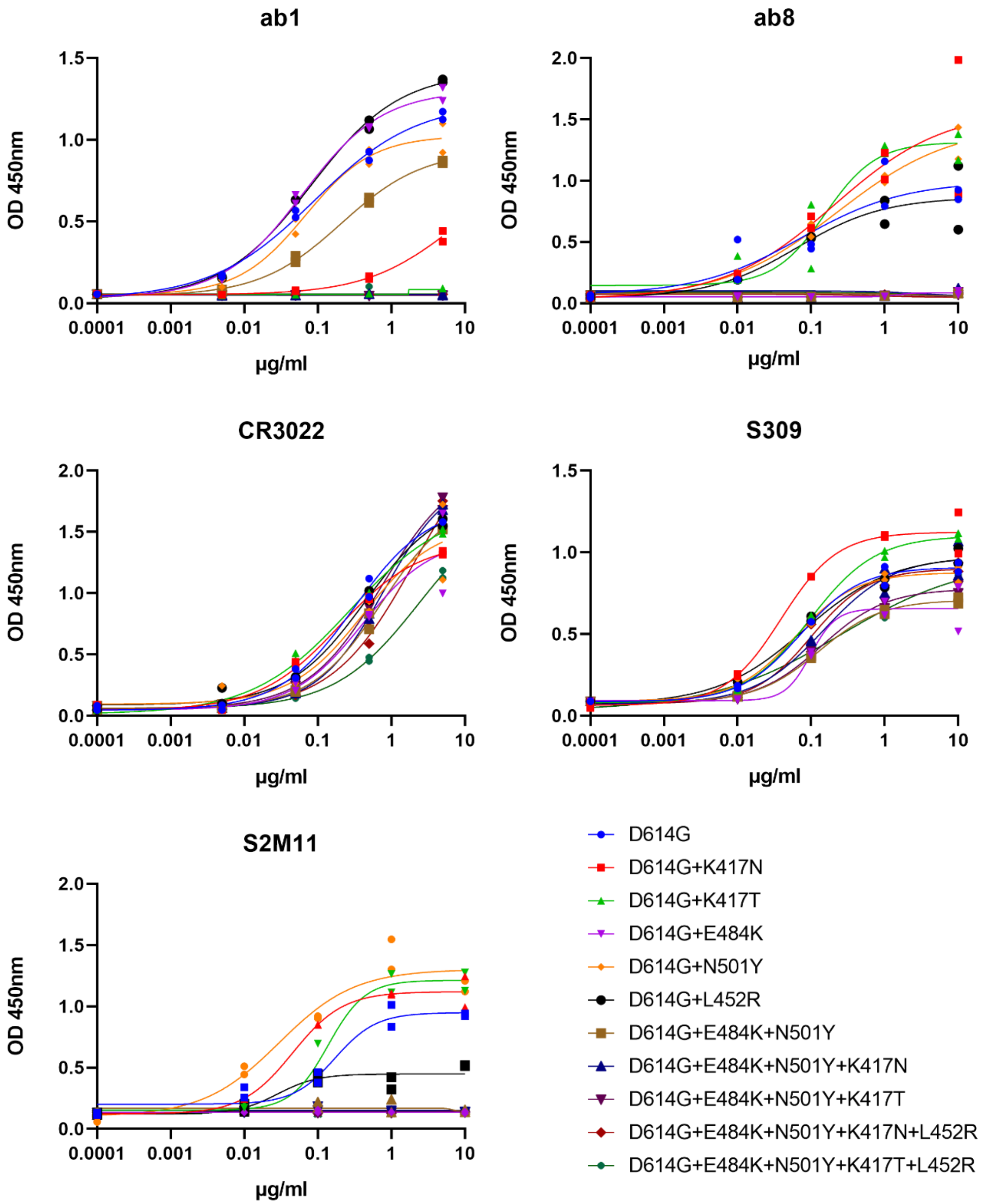


**Figure S19. ELISA analysis of antibody interactions with SARS-CoV-2 VoC RBD mutations, related to figures 4 and S19.** Experiments were performed in duplicate and results are plotted as points.

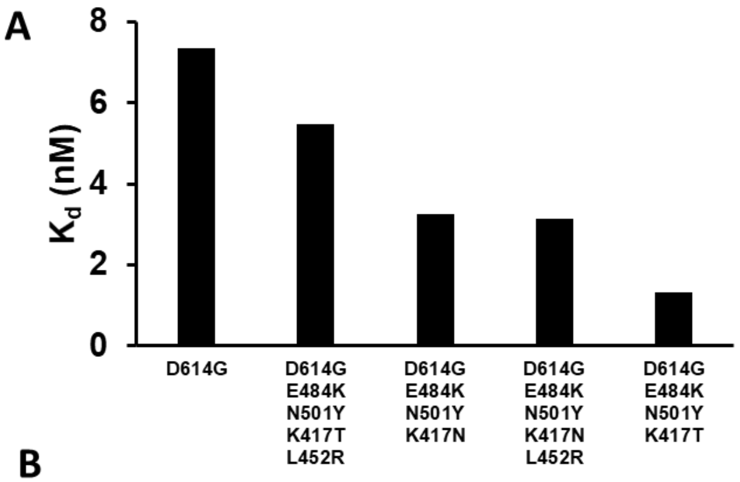


**B**

| Fold Change in S Protein - ACE2 binding relative to D614G Alone | Mutations | Variant of Concern |
|---|---|---|
| 1.00 | D614G | - |
| 1.35 | D614G + E484K + N501Y + K417T + L452R | P.1 + L452R |
| 2.26 | D614G + E484K + N501Y + K417N | P.1.351 |
| 2.35 | D614G + E484K + N501Y + K417N + L452R | P.1.351 + L452R |
| 5.58 | D614G + E484K + N501Y + K417T | P.1 |

<1 Fold Change (Lower affinity)
No Fold Change
>1 Fold Change (Higher affinity)

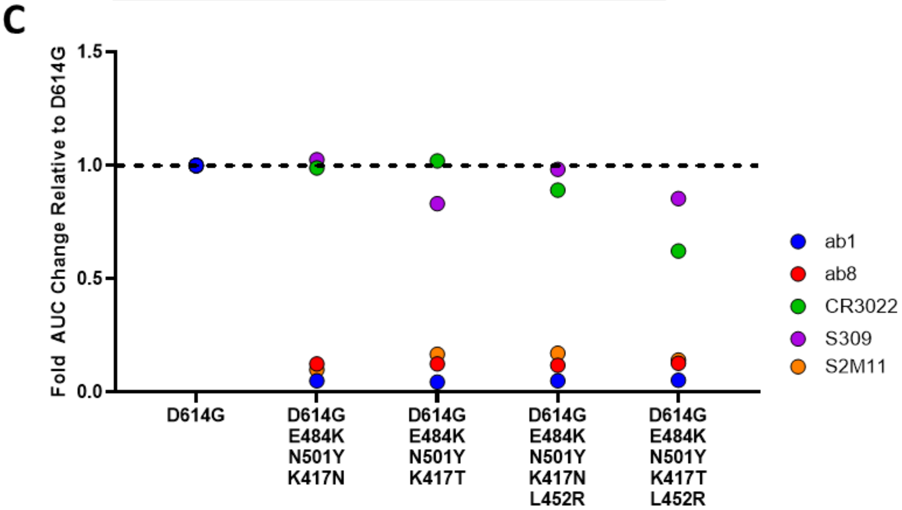


**Figure S20. ACE2 and antibody binding to novel combinatorial VoC RBD mutations. (A)** Affinity (Kd) measurements as measured by biolayer interferometry (BLI). **(B)** Relative fold change differences in S protein - ACE2 affinity (Kd) relative to D614G alone. **(C)** Area under the curve (AUC) fold changes in ELISA binding assays relative to D614G alone for Ab1, Ab8, CR3022, S309, S2M11.

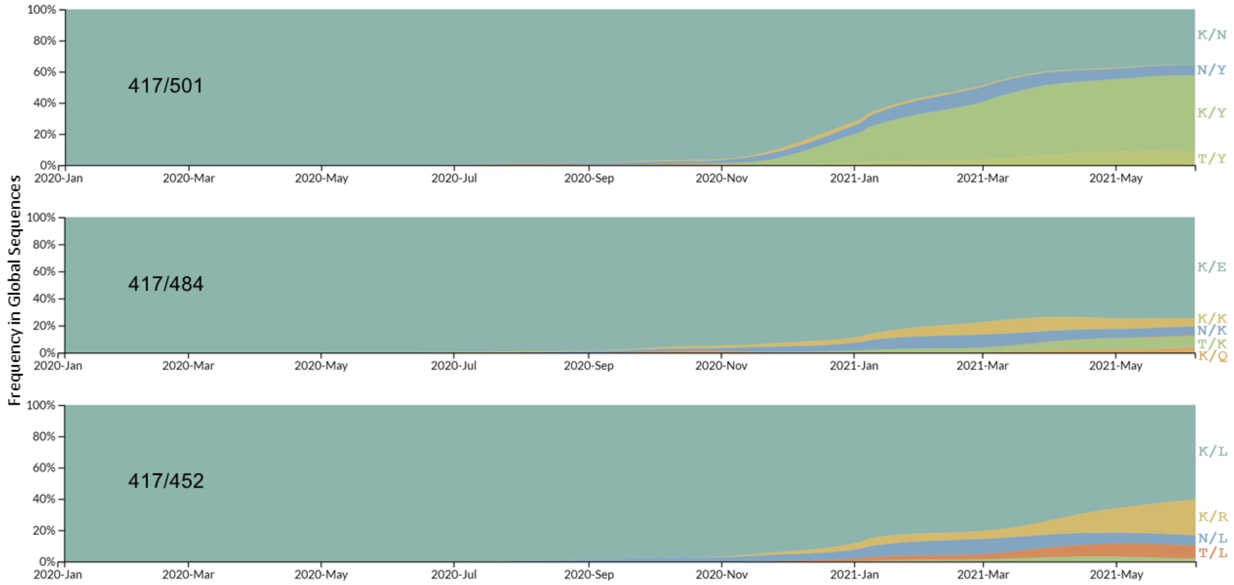


**Figure S21. Analysis of co-mutational prevalence at positions 417, 452, 484, and 501 within the SARS-Cov-2 Spike.** Data from the GISAID sequence databank was analyzed for mutational prevalence between January 2020 and May 2021 at the indicated residue combinations.

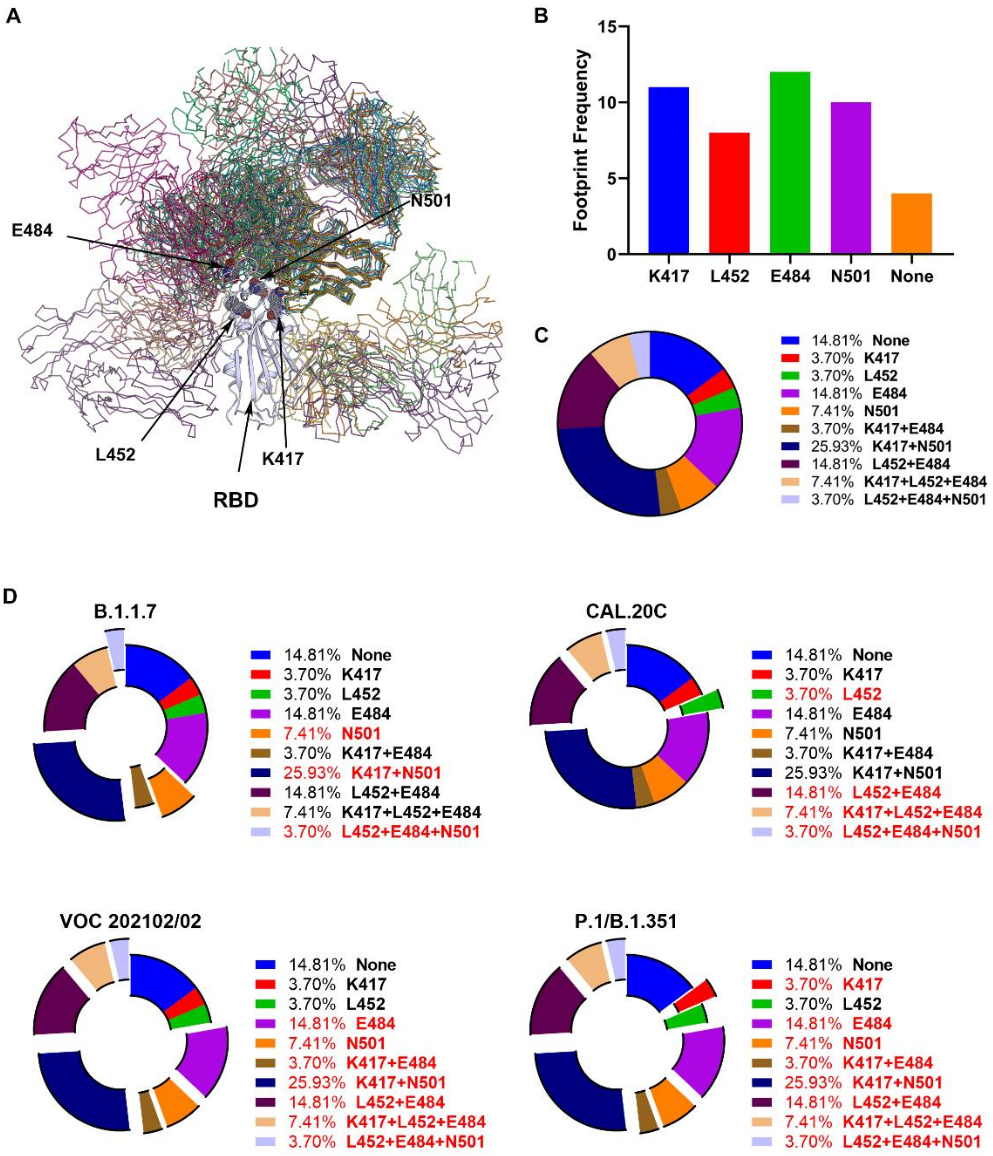


**Figure S22. VoC mutated RBD residues are abundant within the footprints of human derived neutralizing antibodies**. PDB entries of SARS-CoV-2 spike or RBD complexes with antibody fragments isolated from convalescent patients were evaluated for the interaction with RBD positions which are mutated in VoC's. **A)** Structural overlap of all antibodies selected on the SARS-CoV-2 RBD. Mutational positions within the RBD are highlighted. **B)** Frequency of each of the RBD positions which are mutated in VoC's within the footprints of selected antibody-spike/RBD structures. **C)** Proportional analysis of distinct variant RBD positional compositions within the footprints of selected antibody-spike/RBD structures. **D)** Analysis of the overlap between the mutational composition of various VoC's and distinct variant RBD positional compositions within the footprints of selected antibody-spike/RBD structures. Footprints including at least one position mutated within a given VoC are highlighted in red and depicted as slices graphically. Table S1 lists the antibodies and PDB entries selected.

### Table S1. CryoEM data collection and processing parameters, refinement, and validation statistics.

| Structure: | S(D614G) | S(D614G) + ACE2 | | S(D614G,L452R) | S(D614G,L452R) + ACE2 | |
|---|---|---|---|---|---|---|
| | | global refinement | focus refinement | | global refinement | focus refinement |
| | (EMDB 00000) | (EMDB 00000) | (EMDB 00000) | (EMDB 00000) | (EMDB 00000) | (EMDB 00000) |
| | (PDB XXXXX) | (PDB XXXXX) | (PDB XXXXX) | (PDB XXXXX) | (PDB XXXXX) | (PDB XXXXX) |
| **Data collection** | | | | | | |
| Microscope | Titan Krios G4 | Titan Krios G4 | | Titan Krios G4 | Titan Krios G4 | |
| Detector | Falcon4 | Falcon4 | | Falcon4 | Falcon4 | |
| Voltage (kV) | 300 | 300 | | 300 | 300 | |
| Nominal magnification | 155,000 | 155,000 | | 155,000 | 155,000 | |
| Defocus range (µm) | -3.0 to -0.5 | -3.0 to -0.5 | | -3.0 to -0.5 | -3.0 to -0.5 | |
| Physical pixel (Å) | 0.5 | 0.5 | | 0.5 | 0.5 | |
| Electron dose ($e^-/Å^2$) | 40 | 40 | | 40 | 40 | |
| Exposure rate ($e^-/Å^2$/sec) | 24 | 24 | | 24 | 24 | |
| Format of movies | EER | EER | | EER | EER | |
| Number of raw frames | 399 | 399 | | 399 | 399 | |
| Number of movies | 3,726 | 9,153 | | 4,050 | 9,153 | |
| **Data processing** | | | | | | |
| Number of fractions | 40 | 40 | | 40 | 40 | |
| Number of extracted particles | 231,720 | 544,096 | | 292,930 | 732,665 | |
| Number of particles for final map | 120,535 | 65,450 | | 146,886 | 103,023 | |
| Symmetry imposed | C1 | C1 | C1 | C1 | C1 | C1 |
| Resolution (Å) | 3.00 | 2.66 | 2.79 | 2.64 | 2.61 | 3.00 |
| FSC threshold | 0.143 | 0.143 | 0.143 | 0.143 | 0.143 | 0.143 |
| **Refinement** | | | | | | |
| Initial model used | 7MJG | 7MJM,7MJN | 7MJN | 7MJG | 7MJM,7MJN | 7MJN |
| Map sharpening B-factor ($Å^2$) | 76.4 | 54.6 | 80.1 | 63.9 | 60.3 | 94.4 |
| Composition (#) | | | | | | |
| Atoms | 25,236 | 27,000 | 6,557 | 25,245 | 28,597 | 6,553 |
| Residues | 3,120 | 3,318 | 797 | 3,120 | 3,517 | 797 |
| Ligands | NAG:60 | NAG:64 | NAG:7 | NAG:60 | NAG:65 | NAG:7 |
| B-factor ($Å^2$) | | | | | | |
| Protein (min/max/mean) | 69.49/380.50/145.84 | 54.11/368.47/147.56 | 64.46/194.10/102.36 | 58.10/309.37/131.82 | 43.77/400.53/194.53 | 84.28/234.26/131.36 |
| Ligand (min/max/mean) | 92.79/267.84/155.49 | 79.48/283.70/144.37 | 110.92/136.92/121.05 | 77.17/252.41/137.49 | 73.15/403.60/194.71 | 129.89/164.50/147.01 |
| Bonds (RMSD) | | | | | | |
| Length (Å) (# > 4σ) | 0.004 (0) | 0.004 (0) | 0.005 (0) | 0.004 (0) | 0.004 (0) | 0.005 (0) |
| Angles (°) (# > 4σ) | 0.737 (7) | 0.775 (7) | 0.828 (1) | 0.739 (7) | 0.792 (9) | 0.831 (3) |
| CC_mask | 0.84 | 0.83 | 0.87 | 0.84 | 0.83 | 0.83 |
| **Validation** | | | | | | |
| Ramachandran plot | | | | | | |
| Residues favored (%) | 97.14 | 98.35 | 98.74 | 97.20 | 98.04 | 98.61 |
| Residues disallowed (%) | 0.00 | 0.00 | 0.00 | 0.00 | 0.00 | 0.00 |
| Rotamer outliers (%) | 0.00 | 0.00 | 0.00 | 0.00 | 0.00 | 0.00 |
| Clash score | 3.11 | 3.21 | 2.81 | 2.49 | 3.20 | 2.34 |
| MolProbity score | 1.26 | 1.11 | 1.07 | 1.18 | 1.11 | 1.01 |

| Structure: | S(D614G,N501Y) | S(D614G,N501Y) + ACE2 | | S(D614G,N501Y,E484K) | S(D614G,N501Y,E484K) + ACE2 | |
|---|---|---|---|---|---|---|
| | | global refinement | focus refinement | | global refinement | focus refinement |
| | (EMDB 00000) | (EMDB 00000) | (EMDB 00000) | (EMDB 00000) | (EMDB 00000) | (EMDB 00000) |
| | (PDB XXXXX) | (PDB XXXXX) | (PDB XXXXX) | (PDB XXXXX) | (PDB XXXXX) | (PDB XXXXX) |
| **Data collection** | | | | | | |
| Microscope | Titan Krios G4 | Titan Krios G4 | | Titan Krios G4 | Titan Krios G4 | |
| Detector | Falcon4 | Falcon4 | | Falcon4 | Falcon4 | |
| Voltage (kV) | 300 | 300 | | 300 | 300 | |
| Nominal magnification | 155,000 | 155,000 | | 155,000 | 155,000 | |
| Defocus range (µm) | -3.0 to -0.5 | -3.0 to -0.5 | | -3.0 to -0.5 | -3.0 to -0.5 | |
| Physical pixel (Å) | 0.5 | 0.5 | | 0.5 | 0.5 | |
| Electron dose ($e^-/Å^2$) | 40 | 40 | | 40 | 40 | |
| Exposure rate ($e^-/Å^2$/sec) | 24 | 24 | | 24 | 24 | |
| Format of movies | EER | EER | | EER | EER | |
| Number of raw frames | 399 | 399 | | 399 | 399 | |
| Number of movies | 5,112 | 13,955 | | 8,499 | 11,748 | |
| **Data processing** | | | | | | |
| Number of fractions | 40 | 40 | | 40 | 40 | |
| Number of extracted particles | 455,784 | 1,092,783 | | 646,082 | 1,183,976 | |
| Number of particles for final map | 297,629 | 105,418 | 74,520 | 177,775 | 124,036 | |
| Symmetry imposed | C1 | C1 | C1 | C1 | C1 | C1 |
| Resolution (Å) | 2.31 | 2.83 | 3.11 | 3.00 | 2.95 | 3.35 |
| FSC threshold | 0.143 | 0.143 | 0.143 | 0.143 | 0.143 | 0.143 |
| **Refinement** | | | | | | |
| Initial model used | 7MJG | 7MJM,7MJN | 7MJN | 7MJG | 7MJM,7MJN | 7MJN |
| Map sharpening B-factor ($Å^2$) | 52.1 | 65.9 | 95.8 | 87.6 | 76.4 | 118.4 |
| Composition (#) | | | | | | |
| Atoms | 25,248 | 28,599 | 6,554 | 25,248 | 27,029 | 6,554 |
| Residues | 3,120 | 3,517 | 797 | 3,120 | 3,321 | 797 |
| Ligands | NAG:60 | NAG:65 | NAG:7 | NAG:60 | NAG:64 | NAG:7 |
| B-factor ($Å^2$) | | | | | | |
| Protein (min/max/mean) | 46.98/275.84/115.37 | 49.55/433.79/161.85 | 83.07/212.28/125.94 | 68.28/652.80/199.52 | 68.41/427.97/170.67 | 89.97/259.15/152.51 |
| Ligand (min/max/mean) | 64.99/216.96/121.79 | 74.22/355.34/151.18 | 121.16/154.48/139.42 | 94.10/548.41/182.31 | 97.18/347.90/166.43 | 153.20/192.48/172.20 |
| Bonds (RMSD) | | | | | | |
| Length (Å) (# > 4σ) | 0.004 (0) | 0.004 (0) | 0.005 (0) | 0.004 (0) | 0.004 (0) | 0.004 (0) |
| Angles (°) (# > 4σ) | 0.742 (6) | 0.798 (7) | 0.832 (2) | 0.727 (6) | 0.791 (9) | 0.833 (3) |
| CC_mask | 0.83 | 0.82 | 0.85 | 0.80 | 0.82 | 0.84 |
| **Validation** | | | | | | |
| Ramachandran plot | | | | | | |
| Residues favored (%) | 97.20 | 98.01 | 98.61 | 97.17 | 98.04 | 98.36 |
| Residues disallowed (%) | 0.00 | 0.00 | 0.00 | 0.00 | 0.00 | 0.00 |
| Rotamer outliers (%) | 0.00 | 0.03 | 0.14 | 0.00 | 0.00 | 0.00 |
| Clash score | 2.67 | 3.02 | 2.03 | 2.37 | 2.78 | 2.34 |
| MolProbity score | 1.20 | 1.09 | 0.97 | 1.17 | 1.07 | 1.01 |

| Structure: | S(D614G,N501Y,E484K,K417N) | S(D614G,N501Y,E484K,K417N) + ACE2 | | S(D614G,N501Y,E484K,K417T) | S(D614G,N501Y,E484K,K417T) + ACE2 | |
|---|---|---|---|---|---|---|
| | | global refinement | focus refinement | | global refinement | focus refinement |
| | (EMDB 00000) | (EMDB 00000) | (EMDB 00000) | (EMDB 00000) | (EMDB 00000) | (EMDB 00000) |
| | (PDB XXXXX) | (PDB XXXXX) | (PDB XXXXX) | (PDB XXXXX) | (PDB XXXXX) | (PDB XXXXX) |
| **Data collection** | | | | | | |
| Microscope | Titan Krios G4 | Titan Krios G4 | | Titan Krios G4 | Titan Krios G4 | |
| Detector | Falcon4 | Falcon4 | | Falcon4 | Falcon4 | |
| Voltage (kV) | 300 | 300 | | 300 | 300 | |
| Nominal magnification | 155,000 | 155,000 | | 155,000 | 155,000 | |
| Defocus range (µm) | -3.0 to -0.5 | -3.0 to -0.5 | | -3.0 to -0.5 | Falcon4 | |
| Physical pixel (Å) | 0.5 | 0.5 | | 0.5 | 0.5 | |
| Electron dose ($e^-/Å^2$) | 40 | 40 | | 40 | 40 | |
| Exposure rate ($e^-/Å^2$/sec) | 24 | 24 | | 24 | 24 | |
| Format of movies | EER | EER | | EER | EER | |
| Number of raw frames | 399 | 399 | | 399 | 399 | |
| Number of movies | 6,624 | 9,545 | | 4,050 | 20,062 | |
| **Data processing** | | | | | | |
| Number of fractions | 40 | 40 | | 40 | 40 | |
| Number of extracted particles | 479,765 | 996,928 | | 243,776 | 1,486,269 | |
| Number of particles for final map | 241,518 | 127,624 | | 102,603 | 119,936 | |
| Symmetry imposed | C1 | C1 | C1 | C1 | C1 | C1 |
| Resolution (Å) | 2.79 | 2.59 | 2.81 | 2.85 | 2.56 | 3.86 |
| FSC threshold | 0.143 | 0.143 | 0.143 | 0.143 | 0.143 | 0.143 |
| **Refinement** | | | | | | |
| Initial model used | 7MJG | 7MJM,7MJN | 7MJN | 7MJG | 7MJM,7MJN | 7MJN |
| Map sharpening B-factor ($Å^2$) | 76.5 | 60.5 | 85.0 | 61.8 | 69.6 | 99.9 |
| Composition (#) | | | | | | |
| Atoms | 25,245 | 40,083 | 6,553 | 25,242 | 28,595 | 6,552 |
| Residues | 3,120 | 4,908 | 797 | 3,120 | 3,517 | 797 |
| Ligands | NAG:60 | NAG:78 | NAG:7 | NAG:60 | NAG:65 | NAG:7 |
| B-factor ($Å^2$) | | | | | | |
| Protein (min/max/mean) | 65.76/316.41/145.92 | 31.55/367.16/197.13 | 66.20/211.27/109.48 | 60.74/286.20/146.38 | 62.89/370.98/166.16 | 88.65/221.65/129.88 |
| Ligand (min/max/mean) | 86.32/364.95/159.21 | 57.20/382.18/175.49 | 106.84/130.37/120.29 | 84.45/271.98/153.18 | 86.13/310.52/160.15 | 128.85/155.78/144.12 |
| Bonds (RMSD) | | | | | | |
| Length (Å) (# > 4σ) | 0.004 (0) | 0.004 (0) | 0.005 (0) | 0.004 (0) | 0.004 (0) | 0.005 (0) |
| Angles (°) (# > 4σ) | 0.739 (7) | 0.793 (15) | 0.848 (3) | 0.720 (6) | 0.794 (8) | 0.863 (1) |
| CC_mask | 0.78 | 0.77 | 0.86 | 0.80 | 0.83 | 0.85 |
| **Validation** | | | | | | |
| Ramachandran plot | | | | | | |
| Residues favored (%) | 97.27 | 97.86 | 97.73 | 97.23 | 98.01 | 98.61 |
| Residues disallowed (%) | 0.00 | 0.00 | 0.00 | 0.00 | 0.00 | 0.00 |
| Rotamer outliers (%) | 0.00 | 0.00 | 0.14 | 0.00 | 0.03 | 0.00 |
| Clash score | 2.53 | 3.03 | 2.58 | 2.27 | 3.44 | 2.74 |
| MolProbity score | 1.18 | 1.13 | 1.10 | 1.15 | 1.14 | 1.06 |

**Table S2. Antibodies and PDB entries for antibody-RBD complexes selected for analysis in figure S21.**

| Antibody | PDB | References |
|---|---|---|
| CB6 | 7C01 | (Shi et al., 2020) |
| b38 | 7BZ5 | (Wu et al., 2020b) |
| P2B-2F6 Fab | 7BWJ | (Ju et al., 2020) |
| EY6A | 6ZER | (Zhou et al., 2020) |
| COVA2-39 | 7JMP | (Wu et al., 2020a) |
| COVA2-04 | 7JMO | (Wu et al., 2020a) |
| CC12.1 | 6XC2 | (Yuan et al., 2020) |
| CC12.3 | 6XC4 | (Yuan et al., 2020) |
| CV30 | 6XE1 | (Hurlburt et al., 2020) |
| fab 2-4 | 6XEY | (Liu et al., 2020) |
| S2H13 | 7JV2 | (Piccoli et al., 2020) |
| S2A4 | 7JVA | (Piccoli et al., 2020) |
| S2H14 | 7JX3 | (Piccoli et al., 2020) |
| S2X35 | 7JX3 | (Piccoli et al., 2020) |
| 910-30 | 7KS9 | (Banach et al., 2021) |
| C102 | 7K8M | (Barnes et al., 2020a) |
| C105 | 6XCM | (Barnes et al., 2020b) |
| C144 | 7K90 | (Barnes et al., 2020a) |
| C121 | 7K8X | (Barnes et al., 2020a) |
| C002 | 7K8S | (Barnes et al., 2020a) |
| C135 | 7K8Z | (Barnes et al., 2020a) |
| C110 | 7K8V | (Barnes et al., 2020a) |
| C104 | 7K8U | (Barnes et al., 2020a) |
| C119 | 7K8W | (Barnes et al., 2020a) |
| bd23 | 7BYR | (Cao et al., 2020) |
| COVA2-39 | 7JMP | (Wu et al., 2020a) |
| CT-P59 | 7CM4 | (Kim et al., 2021) |

**Table S3. Antibody class definitions and categorization of the antibodies in the present study.**

| Antibody Class | Antibody Class Description | Antibodies in the Present Study |
|---|---|---|
| 1 | Neutralizing antibodies that block ACE2 and bind only to ‘up’ RBDs. | ab1 |
| 2 | ACE2-blocking neutralizing antibodies that bind both up and ‘down’ RBDs and can contact adjacent RBDs. | ab8 |
| 3 | Neutralizing antibodies that bind outside the ACE2 site and recognize both up and down RBDs. | S309 |
| 4 | Previously described antibodies that do not block ACE2 and bind only to up RBDs. | CR3022 |
